# Cocaine exacerbates neurological impairments and neuropathologies in the iTat model of HIV-associated neurocognitive disorder through genome-wide alterations of DNA methylation and gene expression

**DOI:** 10.1101/2021.11.22.469603

**Authors:** Xiaojie Zhao, Fan Zhang, Suresh R. Kandel, Frédéric Brau, Johnny J. He

**Affiliations:** Department of Microbiology and Immunology, Chicago Medical School; Center for Cancer Cell Biology, Immunology and Infection, Rosalind Franklin University, North Chicago, IL 60064; Department of Family Medicine, University of North Texas Health Science Center, Fort Worth, TX, USA; Université Côte d’Azur, CNRS, IPMC, Sophia-Antipolis, 06560, France; School of Graduate and Postdoctoral Studies, Rosalind Franklin University, North Chicago, IL 60064.

**Keywords:** Cocaine, Tat, epigenetic changes, DNA methylation, biological pathways/processes, iTat, HAND

## Abstract

HIV infection of the central nervous system causes HIV-associated neurocognitive disease (HAND) in up to 50% HIV-infected individuals. Cocaine use is prevalent in the HIV-infected population and has been shown to facilitate the HAND progression. However, the cellular and molecular mechanism of the cocaine-facilitated HAND progression remains largely unknown. In this study, we took advantage of the doxycycline inducible and brain-specific HIV Tat transgenic mouse model (iTat) of HAND and characterized effects of chronic cocaine exposure and long-term Tat expression on HAND-associated neurology and neuropathology. We found that cocaine exposure worsened the learning and memory of iTat mice, coupled with dendritic spine swelling, increased synaptophysin expression, and diminished microglia and astrocyte activation. We then employed the single-base resolution whole genome bisulfate sequencing and RNA sequencing and identified 14,838 hypermethylated CpG-related differentially methylated regions (DMR) and 15,800 hypomethylated CpG-related DMR that were linked to 52 down- and 127 up-regulated genes by cocaine and Tat. We further uncovered these genes to be mostly enriched at neuronal function- and cell morphology- and synapse formation-related *ECM-receptor interaction* pathway, and to be linked to behavioral and pathological changes altered by cocaine and Tat. Eight mostly affected genes included four in microglia *Ift172*, *Eif2ak4*, *Pik3c2a,* and *Phf8*, two in astrocytes *Garem1* and *Adgrb3*, and two in neurons *Dcun1d4* and *Adgrb3*. These findings demonstrated for the first time that cocaine and Tat interactively contributed to HAND neurology and neuropathology through genome-wide changes of DNA methylation and gene expression and suggest that targeting epigenetic changes serves as a potentially new therapeutic strategy to treat cocaine use disorder in people living with HAND.

## INTRODUCTION

HIV infection of the central nervous system (CNS) occurs in a majority of HIV-infected individuals and causes HIV-associated neurocognitive disorder (HAND) in up to 50% of the infected population [1–3]. At the cellular level, the primary cell targets for HIV infection are macrophages/microglia and, to a lesser extent, astrocytes [4–9], while neurons that are mostly affected in the brain of HIV-infected individuals are rarely infected. Therefore, a number of indirect mechanisms have been proposed for HAND. One of the major indirect mechanisms is HIV viral protein Tat. Tat is secreted from HIV-infected microglia/microphages and astrocytes [10–12] and taken up by neurons [13–16]. It is detected in the CNS of HIV-infected individuals [17, 18]. Of even more importance is that Tat continues to be expressed in HIV-infected individuals whose HIV replication is effectively suppressed by combination antiretroviral therapy (cART) [17, 19–23]. Recombinant Tat protein, when given *in vivo*, and Tat expression alone in the doxycycline (Dox)-inducible brain-specific HIV Tat transgenic mice (iTat) results in neurological and neuropathological changes reminiscent of those noted in the HIV-infected brain [24–27]. The controllability of the time and duration of Tat expression in this model offers an excellent opportunity to investigate long-term effects of Tat in the context of HIV infection treated with cART and comorbidities such as chronic substance use disorders.

Cocaine use is prevalent in HIV-infected population. 5%-34% HIV-infected population has a history of cocaine use, compared to 0.7%-2% in the general population, and out of approximately 2.2 million regular users of cocaine in the United States, one million individuals develop cocaine use disorder [28–30]. Cocaine impairs the cellular functions and promotes HIV replication [31–33] and disrupts the integrity of the blood brain barrier [34–36]. Cocaine also induces up-regulation of pro-inflammatory mediators and neuroinflammation [37, 38] and facilitates the progression of HAND [39–41]. Furthermore, cocaine causes epigenetic changes. Cocaine alters histone modifications through HDAC, sirtuin, and G9a and transcriptional regulation of genes such as FosB, CDK5, and BNDF at the chromatin level, which contributes to the development and maintenance of addiction [42–44]. Cocaine up-regulates miR-212, leading to an amplification of the stimulatory effects of cocaine on CREB signaling [45]. On the other hand, HIV infection is known to result in genome-wide gene expression, DNA methylation, and miRNA expression. Of particularly note are significant changes of DNA methylation in the HIV-infected brain and Tat-expressing brain, which is linked to accelerated biological aging [46, 47]. Chromatin remodeling is one of major mechanisms of HIV latency and has been exploited as a HIV cure strategy [48]. A number of studies have documented interactive effects between cocaine and Tat. Tat potentiates the psychostimulant effects of cocaine and heightens drug award [27, 49–56]. Tat and cocaine alter metabolism [57–60] and increase the blood brain barrier permeability [61–63]. However, these studies are performed using recombinant Tat protein *in vitro* or short-term Tat expression *in vivo*, and little is not known about whether chronic cocaine use results in epigenetic changes in the context of HIV infection/Tat expression and whether these changes contribute to HAND neurology and neuropathology.

In the study, we determined effects of chronic cocaine exposure and long-term Tat expression on neurology and neuropathology and epigenetic changes. We fed iTat mice with Dox to induce Tat expression for 5 and 11 months, exposed these mice to cocaine for two weeks, kept them drug free for 10 day, and performed behavioral assessments. At the end of behavioral tests, we euthanized the animals and collected brain tissues to perform neuropathology, to isolate genomic DNA for single-base resolution whole genome bisulfate sequencing, or isolate RNA for RNA sequencing. We then used system biology to characterize the gene expression, and biological pathways and processes altered by cocaine and Tat as well as the relationship among epigenetic change-associated gene expression, and biological pathways and processes, neurology, and neuropathology.

## MATERIALS AND METHODS

### Mice and cocaine administration

Wild-type (Wt, C57BL/6) mice were purchased from Jackson Laboratory (Bar Harbor, ME), and iTat mice were created as we described before [26]. All the animal procedures were approved by the Institutional Animal Care and Use Committee. Mice were housed with a 12-hour light and 12-hour dark photoperiod and provided water and food *ad libitum*. Mice were fed with Dox food pellets (0.625g/kg, Envigo, catalog # TD.01306) beginning at day 21 following their weaning and continued for 5 or 11 months, then *i.p.* injected with cocaine (Coc, 30 mg/kg/day, stock: 3 mg/ml) or its solvent saline (Sal) for 14 days. These mice remained on Dox food pellets throughout the Coc injection and behavioral assessments until they were euthanized. Eventually, all mice were fed for a total of either 6.5- or 12.5-month, which were designated as 6m or 12m throughout the study for simplicity.

### Behavioral tests

10 days after final injection (drug free/cessation period), All mice remained drug free for 10 days and then subjected to before behavioral battery tests in the order of increasing stress. ***Elevated plus maze (EPM) test***: In the low light intensity environment, mouse was placed in the central area of EPM apparatus (San Diego Instruments, Part # 7001-0336), faced to open arm, and allowed travel freely between open arms and close arms for five minutes. Then, their travel distance, staying time, and entries in each area/arm were monitored by an infrared camera, and Open Arm Entries, Open Arm Time, and Open Arm Distance were finally used for measuring the anxious status. ***Open field test (OPT)***: Mice were allowed to move freely in an acrylic chamber (San Diego Instruments, Part # 7001-0354) for 10 min, and the Total Distance, Maximum Speed, Central Distance (in the middle square area with 20 cm x 20 cm size), and Central Entries during the movement were recorded for analyzing its locomotor activity and anxious status. ***Rotarod test (RT)***: There are two sessions in two days for RT. Sensitive speed screening session (Day 1): In this session, there were total four trials for each mouse, and the interval between each trial was around 15 min, but no more than 20 min. Mice were first placed on the stationary rod of IITC Rotarod (San Diego Instruments, Part # 2360- 0143) for 30 sec, then started with the acceleration mode from 4 to 45 rpm within 4 min. The speed at which the mouse fell from rod was record, and the average falling-down speed for four trails was calculated. We noted that most young mice fell at 20-35 rpm, and most old mice at 10-20 rpm. So, 30 rpm and 15 rpm were chosen as the sensitive speed for young and old mice, respectively. For the second session test (Day 2), there were two trials for each mouse, and every trial had 4 min test with a fixed sensitive speed (fixed mode). Similarly, mice were placed on the stationary rod for 30 sec first, then started with their sensitive speed. The latency to fall from the rod was record. ***Morris water maze test (MWM)***: The apparatus and protocol were as previously described [64], except that we added to another probe test seven days after first probe test to measure the long-term memory. Briefly, there were two stages including 5-day training and two probe tests. Training stage consisted of four trials. In each trial, mice were put into one quadrant and allowed to freely seek the platform within 90 sec. If they found the platform within 90 sec, they would be allowed to stay on the platform for another 10 sec for memorizing purpose. However, if they failed, they would be put onto the platform to stay for 15 sec. Immobile or floating Mice were excluded from the experiments. First probe test was implemented on the next day after the 5-day training stage with a 60 sec trial, and the second probe test was the same as the first one, but was carried out seven days later. The platform was removed during the probe tests. One day after second probe test, mice were euthanized and the brains were collected. All behavioral tests were record and analyzed by a computerized video tracking system (Anymaze, Stoelting).

### Golgi-Cox staining

After the mice were euthanized, the brain was extracted and sagittally dissected into two hemispheres. Golgi-Cox staining was performed as described [65]. Briefly, one hemisphere was fixed in the Golgi-Cox solution at room temperature for 24 hr and in the fresh Golgi-Cox solution for 10 more days. The tissue was then dehydrated and preserved in tissue-protectant solution at 4°C for 24 hr and in the fresh tissue-protectant solution for seven more days. All these procedures were performed in dark. The tissue was then sagittally sectioned on a virbatome (100 μm thick, Leica, VT1000S). The sections were developed in ammonia solution (3:1) and 5% sodium thiosulfate solution, dehydrated in gradient ethanol and then in xylene, and mounted. All dendritic spine stack images were taken using a Nikon Eclipse 800 microscope with a 100x oil objective, while a 20x objective was used for neuronal branches stack images.

### 3’-Diaminobenzidine (DAB) staining

DAB staining was performed as we described [64]. Briefly, mice were anesthetized by avertin (tribromoethanol) and transcardially perfused with phosphate-buffered saline (PBS) and then 4% paraformaldehyde (PFA). The brains were dissected out, fixed, dehydrated, embedded, and sagittally sectioned with a cryostat (20 μm thick). Floating sections were permeabilized, blocked, probed by Iba-1 antibody (Wako, catalog # 019-19741, and 1:800 dilution) or GFAP antibody (DAKO, catalog # z0334, and 1:500 dilution), inactivated endogenous peroxidases, probed again by a goat anti-rabbit secondary antibody (Southern Biotech, catalog # 4030-05, and 1:200 dilution), and developed using a DAB kit (Abcam, catalog # ab103723). All images were taken using a Nikon Eclipse E800 microscope with a 20x objective for iba-1 staining and 40x for GFAP staining.

### Immunofluorescence staining

Brain sections (20 μm thick) were permeabilized, blocked, probed by NeuN (Millipore-sigma, catalog # MAB377, and 1:500 dilution) and secondary antibody goat anti-mouse 488 (ThermoFisher, catalog # A11001, and 1:500 dilution), and counter stained in 1 μg/ml DAPI. All images were taken using a Nikon Eclipse E800 microscope with a 10x objective.

### Western blotting

Different brain regions, including hippocampus (HIP), caudate putamen (CPU), and cortex (CORT), were dissected out from the fresh frozen brains at -80°C, placed in RIPA buffer (50 mM Tris.HCl, pH 8.0, 280 mM NaCl, 0.5% NP-40, 1% C24H39NaO4, 0.2 mM EDTA, 2 mM EGTA and 10% glycerol) supplemented with protease inhibitors (Millipore-Sigma, catalog# S8830), and briefly sonicated on ice to obtain the lysates. Protein concentrations of the lysates were determined using a Bio-Rad DC protein assay kit (Bio-Rad, catalog # 5000111), the lysates were denatured in the SDS-PAGE loading buffer at 100°C for 10 min, electrophoretically separated by 8-10% SDS-PAGE, transfer onto 0.45 µm polyvinylidene fluoride membrane (GE Healthcare Life Sciences, catalog # 10600023), and probed using appropriate antibodies against PSD-95 (Abcam, catalog # ab18258, and 1:2000 dilution), synaptophysin (SY, Abcam, catalog # ab8049, and 1:1000 dilution), and β-actin (Sigma-Aldrich, catalog # A1978, and 1:2000 dilution). A Bio-Rad ChemicDoc imaging system (Bio-Rad) was used to capture images.

### Image analysis

*For dendritic spine morphology from Golgi staining* All z stack images were firstly projected into 2D images by image J with EDF plugin [66], then input to Imaris software (Bitplane) to analyze Spine Density, Spine Average Area, and Spine Length/Width ratio. All spine were chosen from the terminal branches of neuron dendrites with at least 60 µm in length, every dendritic branch was selected from a different individual neuron, three to four neurons were picked up from each section, three sections were selected from each individual mouse, every experimental group had three mice, and finally 9-12 spines were allocated to each group for statistical analysis. In HIP, only CA regions were included for analysis; in CORT, dendritic branches were randomly chosen from frontal, occipital, and parietal cortex, but every region had at least one dendritic branch; and in CPU, random dendritic branches were selected, as neuron morphology in this region was more uniform. ***For neuronal branches from Golgi staining***: All z stack images were projected into 2D images first, then input to ilastik [67] to run the cell body and branch segmentation, and lastly, analyzed by Cellprofiler [68]. Only the CORT region was selected for this analysis, as no apparent changes were observed in HIP and CPU in our pilot study. ***For microglia morphology from Iba-1 DAB staining***: All branches of microglia were pronounced by ilastik and analyzed by Cellprofiler. ***For astrocyte morphology from GFAP DAB staining***: Image J was used to segment the cell body region and count the cell number; and astrocyte branches were pronounced by ilastik and calculated by Cellprofiler. We only chose HIP dentate gyrus (DG) region in this analysis because of its significance as well as most if not all negative GFAP staining in CORT and CPU regions. ***For neuron number from NeuN immunofluorescence staining***: In HIP, only a specific area located at CA2-CA3 region (framed by white box in **Fig. 2c**) was chosen for analysis; the green (NeuN+) and blue (DAPI+) channels in this area were split by image J, then the cell body signals in green channel were pronounced by ilastik and calculated by Cellprofiler. The blue channel in HIP and all staining signals in CORT and CPU regions were directly processed by Cellprofiler. In HIP and CPU, three random areas were chosen from each section; in CORT, three areas were chosen for each of frontal, occipital and parietal cortex from each section, thus three areas from three cortex regions were averaged for CORT analysis; for all the brain regions, three to five sections were selected from each mouse, every group had three mice, and there were 9-15 sections for each group for statistical analysis.

### DNA and RNA isolation, library preparation, and sequencing

Genomic DNA was isolated from one HIP hemisphere by DNeasy Blood & Tissue Kit (Qiagen, catalog # 69504), the DNA quality was first confirmed by Nanodrop (ThermoFisher, catalog # 13-400-518) with 260/280 ratio at 1.95-2.0 (in 10 mM Tris·Cl, pH 7.5 solution) and then verified by 1% agarose gel analysis to make sure no DNA degradation and RNA contamination. Genomic DNA libraries were prepared genomic DNA (10 ng) using a Pico Methyl-Seq Library Prep Kit (ZYMO, catalog # D5456). MQ beads (Epigentik, catalog # P-1059) were used in place of the column purification system in the library kit for higher DNA recovery. The quality of DNA libraries was determined to be 380-520 bp of the median size and have no contamination of primer dimers. The libraries were sequenced using the Illumina NovaSeq 6000 S4 Paired-end-150 system. RNA was isolated from the other HIP hemisphere of the same mice by TRIzol reagent (ThermoFisher, catalog # 15596026) and further purified using a RNA Clean & Concentrator Kit (ZYMO, catalog # R1013). The RNA was determined to be 9.5-9.9 of RNA integrity number using an Agilent Bioanalyzer 2100. The RNA libraries were prepared from the RNA using a QIAseq Stranded mRNA Select kit (Qiagen, catalog # 180773). Briefly, RNA (800 ng) was used to generate poly- A+ enriched RNA, the concentration of which was determined using a Qubit 4 Fluorometer (ThermoFisher, catalog # Q33238). poly-A+ enriched RNA (10 ng) was used to synthesize the RNA libraries using the kit. The quality of libraries was determined to be 300-500 bp of the median size and have no contamination of primer dimers using a bioanalyzer. The libraries were sequenced using the Illumina HiSeq 4000 Single-end-50 system.

### Sequencing data processing and analysis

A bioinformatics analysis pipeline nf-core/methylseq [69] was used to process the methylation (Bisulfite) sequencing data. Briefly, we input the FASTQ raw data, aligned the reads, and performed extensive quality-control on the results. In the pipeline, FASTQC (v 0.11.9) (http://www.bioinformatics.babraham.ac.uk/projects/fastqc/) was chosen for quality control and Trim Galore! (v0.6.5) (http://www.bioinformatics.babraham.ac.uk/projects/trim_galore/) for adapter sequence trimming with default parameters. Trimmed sequences were then mapped to the Mus musculus assembly (GRCm38/mm10) from Genome Reference Consortium, using Bismark 0.22.3 with the alignment tool Bowtie2 2.4.2. Sequence duplicates were further removed by command “deduplicate_bismark” and context-dependent methylation (CpG and CpH) were extracted by command “bismark_methylation_extractor”.

### Differential methylation calculation

M-value was used as metrics to measure methylation levels and differential analysis of methylation levels, which was defined as the log2 ratio of the intensities of methylated probe versus unmethylated probe as followed [70]:

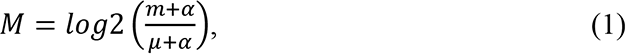

where *m* and *μ* are methylated and unmethylated level, respectively, measured by the methylated and unmethylated probes for an interrogated CpG site respectively, and *α* is a constant offset and set as 1e-3. We modeled each CpG or CpH site once for differential methylation analysis. For each site and preset region, we considered the following linear mixed model to determine differential methylation:

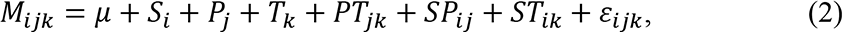

where *ε*_*ijk*_∼*N*(0, *σ*^2^). The fixed effects in equation are *μ* (mean), *S*_*i*_ (sex, male or female), *P*_*j*_ (genotype, iTat or Wt), *T*_*k*_ (treatment, Coc or Sal), *PT*_*jk*_ (genotype-treatment interaction), *SP*_*ij*_ (sex-genotype interaction), and *ST*_*ik*_ (sex-treatment interaction); *i* = 1,2 stands for male and female, respectively; *j* = 1,2 stands for iTat and Wt mice, respectively; and *k* = 1,2 stands for Coc and Sal treatment, respectively. The variance components in the linear mixed model were estimated using the residual maximum likelihood (REML) approach [71]. We used R program (http://www.r-project.org, version 4.0.4) to fit linear mixed models to each site and preset region.

The lsmeans package was used for testing the differences in combinations of levels among genotype and treatment, and for tests of Difference-in-Differences in the interaction between genotype and treatment. After initial analysis, we did not find significant difference by sex factor. We then only considered two factors, genotype and treatment, and defined the vectors with only four values [Wt mice with Sal (Wt-Sal), iTat mice with Sal (iTat-Sal), Wt mice with Coc (Wt-Coc), and iTat mice with Coc (iTat-Coc)] to represent the means we used to compare. Thereafter, we built the custom comparisons via the Contrast () function in lsmeans package. For example, Wt-Coc vs. Wt-Sal is defined as (0, 1, 0, -1), iTat-Sal vs. Wt-Sal as (0, 0, 1, -1), and interaction between type (iTat/Wt) and treatment (Coc/Sal) as (1, -1, -1, 1).

### Annotation with custom regions

Our preset regions were downloaded from annotatr package [72], which provides genomic annotations including 1 to 5 kb, promoters, 3UTRs, 5UTRs, exons, exon boundaries, introns, and intergenic. Then, three critical regions, including promotors, exons, and introns were parsed and merged with GRCm38 gene annotation file for region analysis.

### RNA sequencing (RNA-Seq) data processing

Reads were mapped to GRCm38 reference genomes with hisat2 (v2 2.2.1). Then, SAM files were converted and sorted with smarttools (v1.11). And, featureCounts (v 2.0.1.13) was used to quantify reads. For each exon, we applied the quantified expression value *Y*_*ijk*_ the same linear mixed model as shown in equ (2) to determine differential expression:

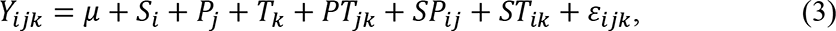

where variables and subscripts are same as equ (2), except for *Y*_*ijk*_ which is a quantified expression value.

### Gene-linked DMR

To determine the potential genes which were possibly regulated by differentially methylated regions (DMR). We defined Gene-linked DMR as the differentially methylated regions mapped to or close to genes that are differentially expressed (DEG) in RNA-Seq. Gene expression values were measured at the exon level from RNA-Seq by feature Counts, in order to increase mapping with DMR. We merged the DMR annotated from annotatr package [72] and differentially expressed exons from RNA-Seq by transcript ID. There were totally four types of gene-linked DMR: Hyper-methylated DMR (Hyper-DMR) linked to up-regulated genes (Up-DEG), Hyper-DMR linked to down-regulated genes (Down-DEG), Hypo-methylated DMR (Hypo-DMR) linked to Up-DEG, and Hypo-DMR linked to Down-DEG. Only Hyper-DMR linked to Down-DEG and Hypo-DMR linked to Up-DEG were used for down-stream analysis.

### KEGG and Gene Ontology analysis

We performed pathway analysis through the Database for Annotation, Visualization and Integrated Discovery (DAVID) v6.8 [73]. Functional Annotation Table for the three factors, Tat, Coc, and Tat X Coc, was downloaded through the DAVID API Server.

### Machine learning for cell type prediction

In order to predict the cell types for gene-linked DMR, we first downloaded the top 6000 ranked cell type-enriched mouse genes from the reference [74] and mapped them with the RNA-Seq transcriptome database of purified cell classes of the brain [75]. The Support Vector Machine [76] was used to build a learning model from the mapped genes, and then the trained model was applied to predict the cell type for the unmapped genes and generated a database with cell type information. Finally, we mapped the gene-linked DMR to this database.

### Statistical analysis

Four-way repeated measures ANOVA was used in MWZ training stages, and all other studies used either two-way or four-way ANOVA, whenever applicable; Bonferroni test was used for all post hoc analyses. All statistical analyses were performed using the IBM SPSS 20; and *p*<0.05 was considered significant and marked as *, # or $ for comparisons among different groups; *p*<0.01 and *p*<0.001 were both considered highly significant and marked as ** and ***, respectively. For all bioinformatic data analysis, R program with *q*<0.05 or *p*<0.05 was used, and other details were provided the sections above. Interactive effect between Tat and cocaine was defined by ANOVA tests or Linear mix model analysis. If there was no interactive effect and combined effect of these two factors was more than each of the factors alone, it was defined as additive effect.

## RESULTS

### Cocaine exposure worsened learning and memory impairments by Tat

To determine effects of chronic cocaine use on HIV-associated neurocognitive disorder in young and middle-aged adults with long-term HIV infection under effective cART suppression, we set up the following experimental scheme (**Fig. 1a**): iTat mice were fed with Dox immediately after weaning for 5 and 11 months, injected cocaine (Coc, 30 mg/kg/day) for 14 days [77–79], kept drug-free for 10 days, and subjected to a battery of behavioral tests. Wt mice and saline (Sal) were included as the controls for iTat mice and Coc, respectively. There were a total of 194 mice in the study, which were randomly assigned (10-16 mice/group) to 16 experimental groups [2 genotypes (Wt, iTat) x 2 sexes (male, female) x 2 age groups (6m, 12m) x 2 treatments (Sal, Coc). Behavioral tests were performed in the order of increasing stress level, and they were elevated plus maze (EPM) for anxiety, open field test (OPT) for locomotor activity and anxiety, rotarod test (RT) for balance and coordination, tail suspension test (TST), forced swim test (FST) for depressive status, and Morris water maze (MWM) for learning and memory.

**Figure 1.**
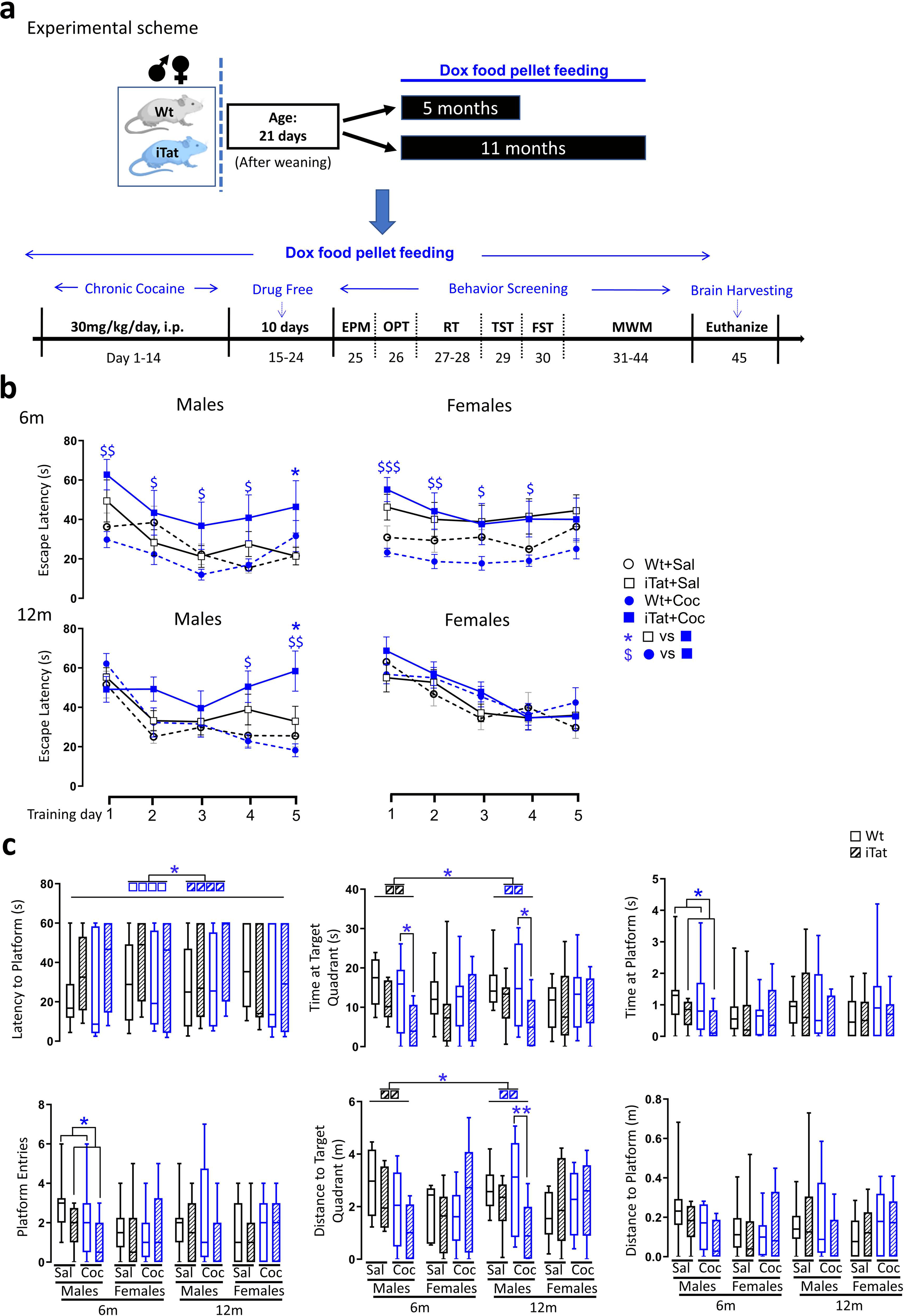
Effects of cocaine on the learning and memory of iTat mice. **a**. Experimental scheme. Wt and iTat mice were fed with Dox food pellets for 5 or 11 months from day 21 when they were weaned, given cocaine i.p. (30mg/kg/day) for 14 day, remained drug free for 10 days, and subjected to behavioral assessments: EPM, OPT, RT, TST, FST, and MWM. Dox food pellets continued throughout the studies. Mice were euthanized to collect tissues for pathology or DNA/RNA isolation following MWM. **b**. Escape latency in the MWZ training stages. **c.** Short-term memory from the first probe test demonstrated by Latency to Platform, Platform Entries, Time at Target Quadrant, Distance to Target Quadrant, Time at Platform, and Distance to Platform. *P*<0.05 was considered significant and marked as * or $ for comparisons among different groups; *p*<0.01 and *p*<0.001 were both considered highly significant and marked as ** and ***, respectively. Sal: saline, Coc: cocaine.

MWM consisted of a 5-day training for leaning ability, measured by Escape Latency and Cumulative Distance, and two probe tests for short-term and long-term memory, measured by Latency to Platform, Platform Entries, Time at Target Quadrant, Distance to Target Quadrant, Time at Platform, and Distance to Platform. During the 5-day training, iTat mice showed longer Escape Latency trend than Wt mice in the absence of cocaine, Coc prolonged Escape Latency in iTat mice while only slightly shortened Escape Latency in Wt mice (**Fig. 1b**). Cumulative Distance showed the same pattern, but with the more pronounced impact of Tat on 6m females (**Suppl. Fig. 1a, Suppl. Table 1**). 12m females had less pronounced effects of Tat and Coc than males of 6m and 12m and 6m females. For the first probe test (short-term memory), which was performed one day after the last training, iTat 6m males showed shorter Time at Platform and fewer Platform Events than Wt 6m males, and Coc showed shorter Latency to Platform in all iTat mice than Wt mice, less Time at Target Quadrant in iTat males than Wt males of both 6m and 12m groups and shorter Distance to Target Quadrant in iTat 12m males than Wt 12m males (**Fig. 1c**). In addition, Coc showed shorter Time at Target Quadrant and shorter Distance to Target Quadrant than Sal in iTat males. For the second probe test, which was performed 7 days after first probe test for long-term memory, iTat mice showed shorter Time at Target Quadrant than Wt in 6m females and 12m males, shorter Time at Platform, fewer Platform Events, shorter Distance to Target Quadrant, and shorter Distance to Platform than Wt in 12m males when given Coc, and iTat showed longer Latency to Platform than Wt in 12m males, regardless of Coc or Sal treatment (**Suppl. Fig. 1b**). In addition, Coc showed longer Distance to Target Quadrant than Sal in iTat males.

For EPM, measured by Open Arm Entries, Open Arm Time, and Open Arm Distance, iTat showed more Open Arm Entries than Wt in 6m males and 6m females, longer Open Arm Time than Wt in 12m males, and longer Open Arm Distance than Wt in 12m males (**Suppl. Fig. 2a, Suppl. Table 2**). Coc showed more Open Arm Entries and longer Open Arm Distance than Sal in iTat 6m, WT 6m females, and iTat 12m of both males and females. For OPT, measured by Total Distance and Maximum Speed for locomotor activity, and Central Distance and Central Entries for anxiety, iTat showed shorter Total Distance than Wt in 12m females, lower Maximum Speed than Wt in 6m females and 12m females, longer Central Distance than Wt in 12m males, and more Central Entries than Wt in 12m males (**Suppl. Fig. 2b**). Coc showed shorter longer Total Distance than Sal in iTat 6m females, higher Maxiumun Speed than Wt in iTat 12m females, longer Central Distance than Sal in iTat 6m females and Wt 12m males, and more Central Entries than Sal in 6m females of both Wt and iTat. For RT, measured by Latency to Fall for balance and coordination ability, iTat showed shorter Latency to Fall than Wt in 6m females at the speed of 30 rpm, Coc showed longer Latency to Fall that Sal in 12m females (**Suppl. Fig. 2c**). For TST and FST, measured by Immobile Time for depressive status, Coc showed shorter Immobile Time of TST than Sal in iTat 12 m females (**Suppl. Fig. 2d**) and shorter Immobile Time of FST than Sal in Wt 12m females (**Suppl. Fig. 2e**). In addition, iTat showed shorter Immobile Time of FST in 6m males than Wt, regardless of Coc or Sal treatment. Taken together, these results demonstrated that iTat mice exhibited impaired learning and memory and Coc worsened impaired learning and memory of iTat mice. These results also showed that iTat mice exhibited impaired locomotor activity and balance and coordination, sex- and age-dependent higher threshold of anxiety, and somewhat depressive-like behaviors, but Coc improved these behaviors of iTat mice.

### Cocaine exposure led to dendritic spine swelling without altering the number of neurons and neuronal branches in the presence of Tat

We then investigated changes of three major brain cells: neurons, microglia, and astrocytes of iTat mice exposed to cocaine. We first performed the Golgi-Cox staining to determine morphological changes in neurons in hippocampus (HIP), cortex (CORT), and caudate putamen of dorsal striatum (CPU). Neurons were characterized by Spine Density, Average Area, and Length/Width ratio and quantitated by projecting stack images and modeling the spines.

In HIP (**Fig. 2a**), iTat showed lower Spine Density than Wt in all males. However, Coc showed higher Spine Density than Sal in all mice, and the effect was more pronounced in iTat 6m females than Wt (**left panel, Fig. 2b**). Coc showed higher Spine Average Area than Sal in Wt 6m males, iTat had smaller Spine Average Area than Wt in 6m females and higher Spine Average Area than Wt in 12m of both males and females exposed to Coc (**middle panel, Fig. 2b**). Coc showed higher Spine Length/Width ratio than Sal in Wt 6m males, and iTat showed lower Spine Length/Width ratio than Wt in 6m females (**right panel, Fig. 2b**). We also performed the immunofluorescent staining against NeuN to determine the number of neurons (**Fig. 2c**) using the number of DAPI+ stained cells as a reference. iTat showed lower NeuN+ cells/DAPI+ cells than Wt in CA2 and CA3 regions (**bracketed, Fig. 2c**) of all males and females of both 6m and 12m (**Fig. 2d**). Similar trends of changes in neuron morphology and the number of neurons were obtained in CORT (**Suppl. Fig. 3, Suppl. Table 3**) and CPU (**Suppl. Fig. 4, Suppl. Table 3**). Some notable differences included that generally larger dendritic spines in CORT and CPU made the differences more pronounced, that Tat-associated sparse and smaller dendritic spines in CORT was absent and to a lesser extent in CPU, and that even much thinner Tat-associated dendritic spines were found in CPU. Furthermore, we determined the length of neuronal branches including dendrites and axons and quantitated by projecting stack images and segmenting the cell body (**Fig. 3a**). In CORT, only iTat showed shorter Branch Length than Wt in 6m males (**Fig. 3b**). No significant differences in Branch Length were noted in HIP and CPU among all groups (**data not shown**). These results together showed that iTat showed lower density of dendritic spines with shorter, wider and smaller size morphology similar to the stubby shape [80], and Coc showed higher density with longer and thinner morphology similar to the filopodia or thin type [80], whereas iTat exposed to Coc showed higher density with shorter and bolder morphology close to the mushroom type but with a much larger size, indicating that cocaine exposure leads to dysgenesis-like dendritic spine swelling or synaptopathology.

**Figure 2.**
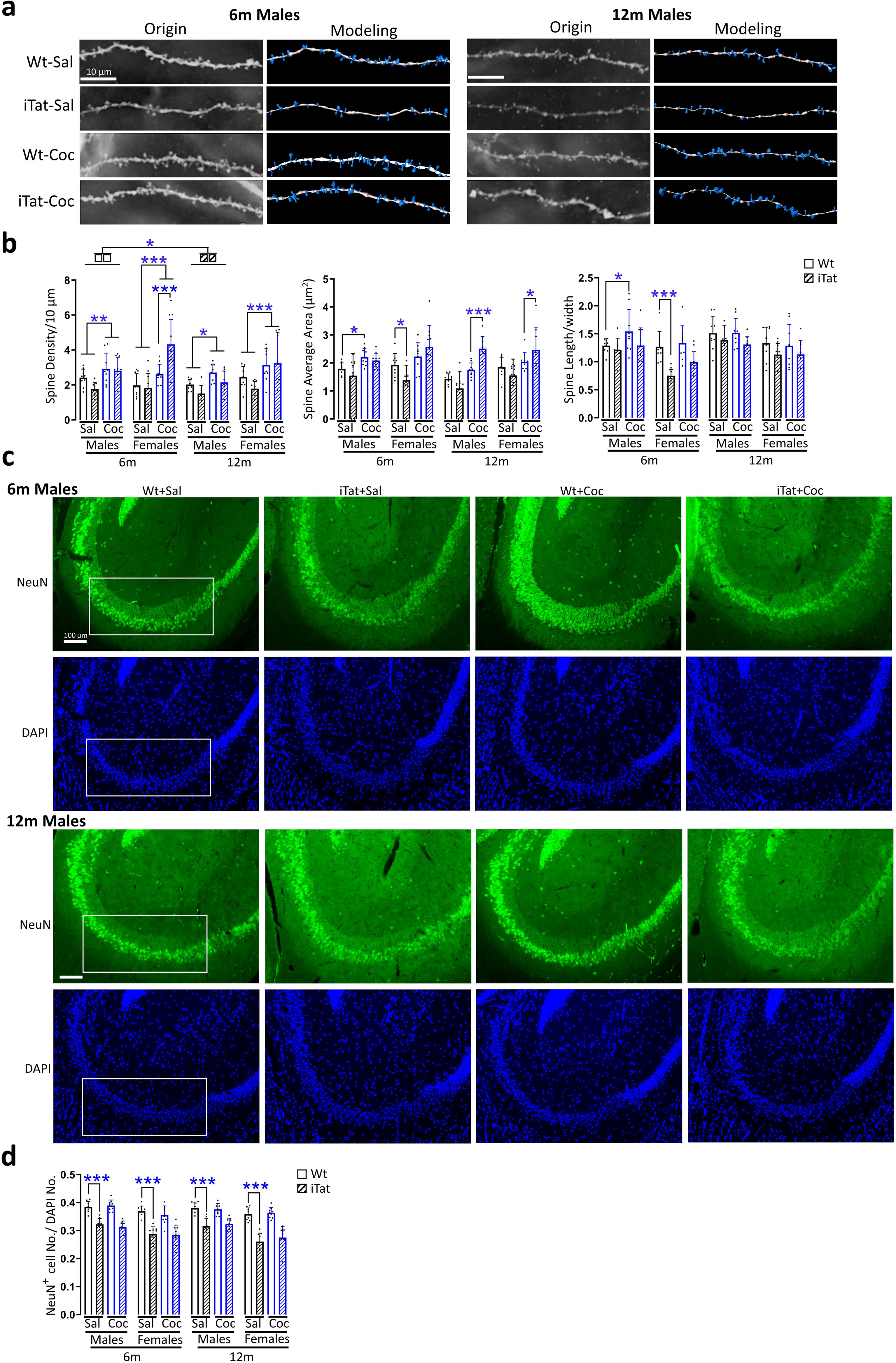
Effects of cocaine dendritic spine morphology and the number of neurons in HIP of iTat mice. **a** & **b**. Mouse brain sections were stained in the Golgi-Cox solution (**a**), dendritic spines in CA1-4 regions were modeled and quantified by Imaris for spine density, occupied area and length/width (**b**). **c** & **d**. Mouse brain sections were immune stained for NeuN and counter stained in DAPI (**c**), the total number of neurons (NeuN-positive) in CA2-3 regions (marked by white line boxes) was quantified and expressed as a fraction of the total number of cells (DAPI-positive) (**d**). Representative images (**a** & **c**) were chosen from 6- and 12-months male mice groups. *P*<0.05 was considered significant and marked as *, and *p*<0.01 and *p*<0.001 were both considered highly significant and marked as ** and ***, respectively. Scale bars: 10 µm for **a**, 100 µm for **c**. Sal: saline, Coc: cocaine.

**Figure 3.**
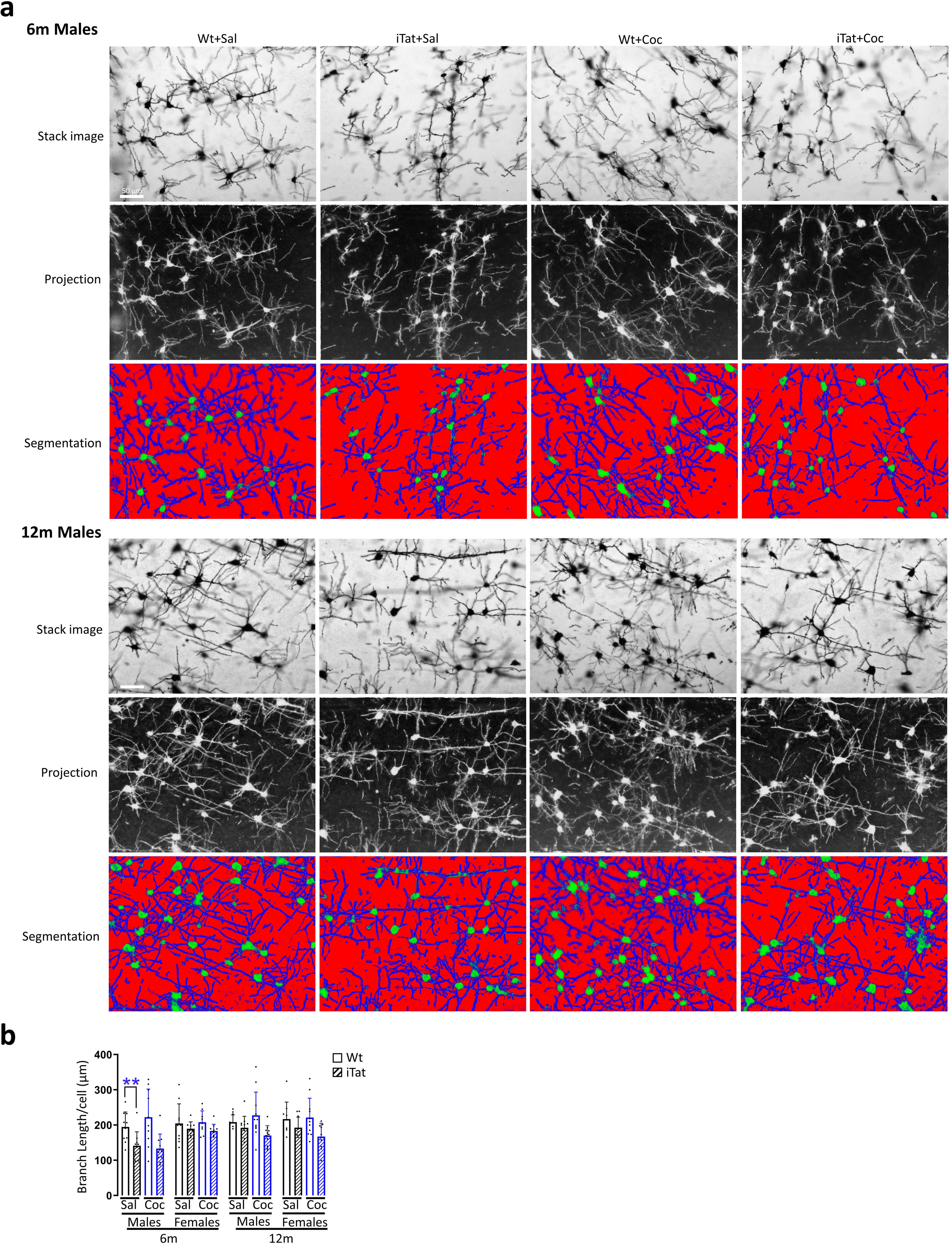
Effects of cocaine on dendrites and axons of neurons in CORT of iTat mice. Mouse brain sections were stained in the Golgi-Cox solution. z stack images of dendrites and axons of neurons in CORT were taken, projected in the same layer by Fiji, and segmented out all branches by ilastik (**a**). Representative images were chosen from parietal cortex in 6- and 12- months male mice groups. The branch length in each individual cell was calculated by Cellprofiler (**b)**. *p*<0.05 was considered significant and marked as *, and *p*<0.01 and *p*<0.001 were both considered highly significant and marked as ** and ***, respectively. Scale bars in **a**: 50 µm. Sal: saline, Coc: cocaine.

### Cocaine exposure led to significant increases of synaptophysin expression in the presence of Tat

We next performed Western blotting to determine expression of two important neuronal functional markers post-synaptic postsynaptic density protein 95 (PSD-95) and pre-synaptic synaptophysin (SYP) in the brain of these mice (**Fig. 4a**). In HIP, Coc showed lower PSD-95 expression than Sal in all groups, and iTat showed higher PSD-95 expression than Wt in all Coc groups (**Fig. 4b**). Coc showed significant higher SYP expression than Sal in all groups except for 6m females, and iTat showed higher SYP expression than Wt in all Coc groups. Similar trends were noted in both CORT (**Suppl. Fig. 5**) and CPU (**Suppl. Fig. 6**), although the differences were smaller in these two regions compared to HIP and iTat showed higher PSD-95 and SYP than Wt in 12m females. One consistent finding from all three brain regions was that Coc increased considerably more SYP expression in iTat mice than Wt in all groups.

**Figure 4.**
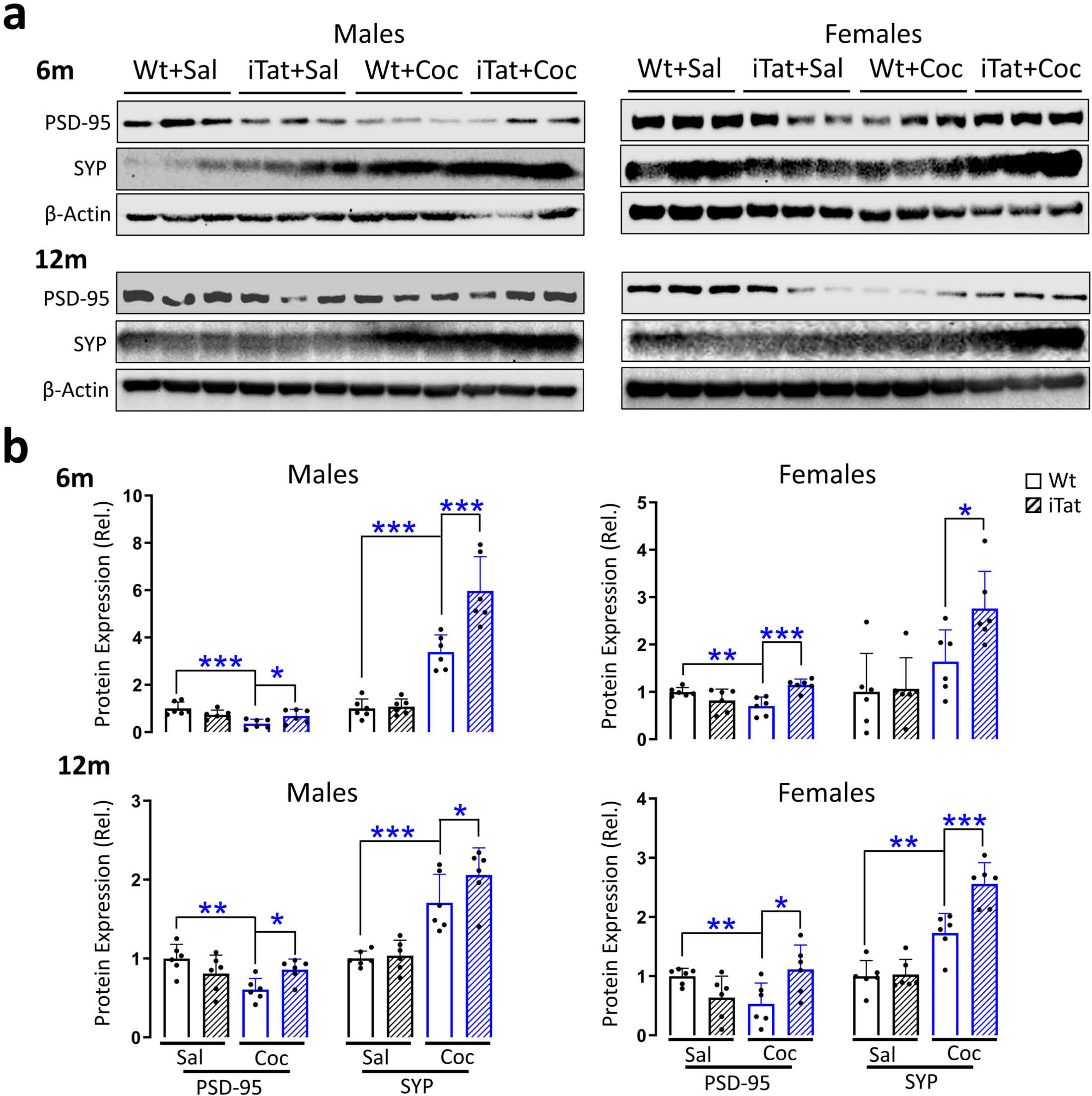
Effects of cocaine on synaptophysin expression in hippocampus (HIP) of iTat mice. HIP brain regions were dissected from the mouse brain and processed for lysates. SYP and PSD-95 expression in the lysates were determined by Western blotting (**a**), quantified by Fiji, normalized to the loading control β-actin, and calculated using the Wt+Sal as a reference, which was set at 1 (**b**). *p*<0.05 was considered significant and marked as *, and *p*<0.01 and *p*<0.001 were both considered highly significant and marked as ** and ***, respectively. Sal: saline, Coc: cocaine.

### Cocaine exposure diminished microglia activation by Tat

We then performed immunohistochemistry staining for microglia marker Iba-1 and determined if there were changes in the number and morphology of microglia. The cell body and branches of Iba-1+ cells were segmented, and the branches were skeletonized. The branch length and endpoints were quantified for each individual cell. In HIP (**Fig. 5a**), iTat had more microglia than Wt in all groups except for 12m females (**left panel, Fig. 5b**), and longer Branch Length (**middle panel, Fig. 5b**) and more Branch Endpoints (**right panel, Fig. 5b**) than Wt in all groups. Coc had longer Branch Length and more Branch Endpoints than Sal in 6m of both males and female, while Coc had shorter Branch Length and fewer Branch Endpoints than Sal in iTat 6m of both males and females and iTat 12m males. The same findings were noted in CORT (**Suppl. Fig. 7, Suppl. Table 4**) and CPU (**Suppl. Fig. 8, Suppl. Table 4**), only differing in statistical significances. These results showed that Tat activated microglia to proliferate along with longer branches and more endpoints, Coc did not cause microglia to proliferate but with longer branches and more endpoints, but unexpectedly, Coc and Tat led to decreased branch length and endpoints of microglia while maintained the higher number of microglia.

**Figure 5.**
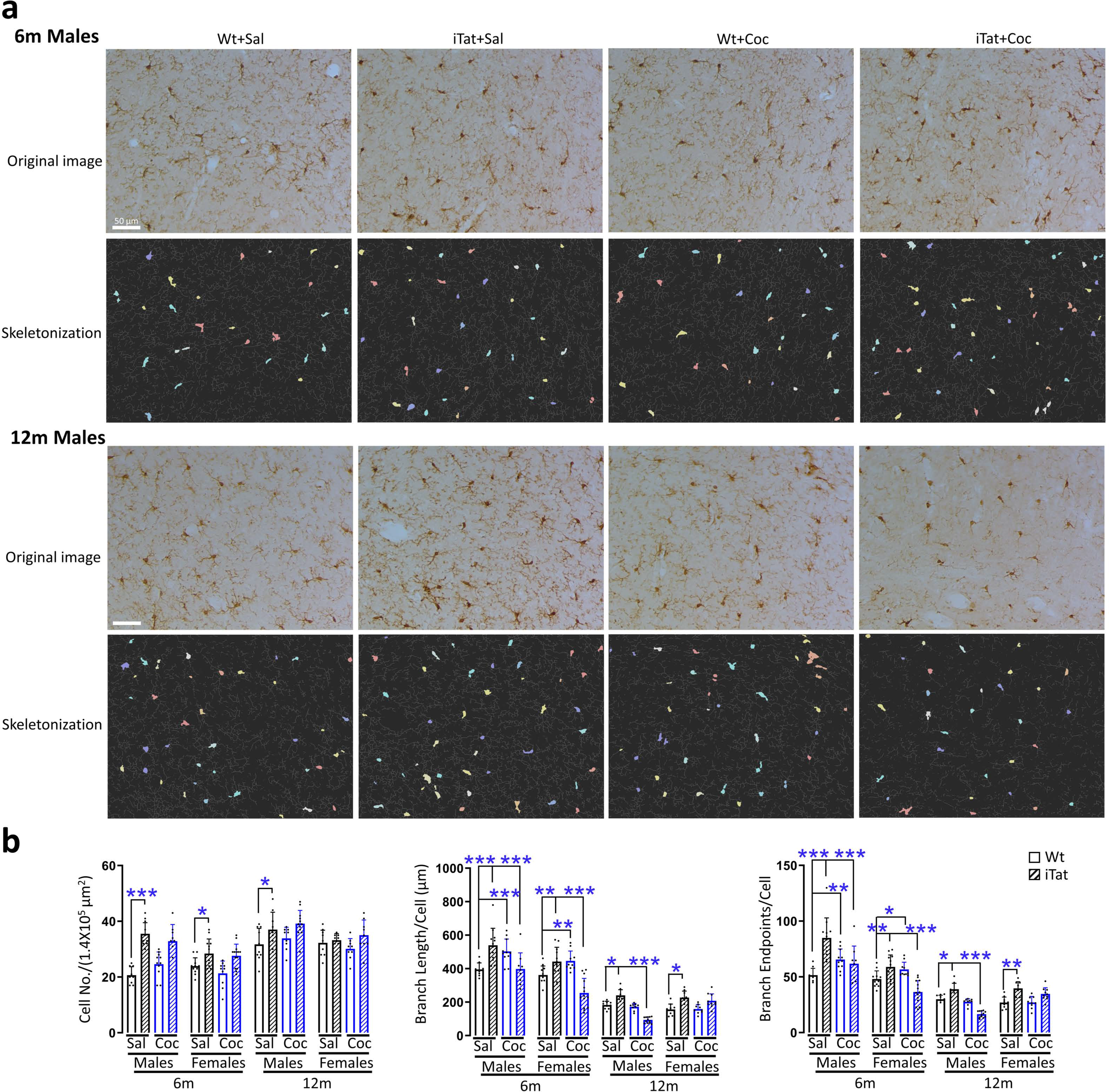
Effects of cocaine on microglia of HIP of iTat mice. Mouse brain sections were stained for Iba-1, the cell bodies of microglia were segmented and the branches were skeletonized by Cellprofiler **(a**). The total number of microglia in every view field (1.4 x 10^5^ μm^2^), and the branch length of microglia and the number of the ending points in each microglia were further calculated by Cellprofiler **(b**). Representative images were chosen from 6- and 12- months male mice groups. *P*<0.05 was considered significant and marked as *, and *p*<0.01 and *p*<0.001 were both considered highly significant and marked as ** and ***, respectively. Scale bars in **a**: 50 µm. Sal: saline, Coc: cocaine.

### Cocaine exposure diminished astrocyte activation by Tat

We also performed immunohistochemistry staining for astrocyte marker glial fibrillary acidic protein (GFAP) and characterized the morphological changes of astrocytes in these mice. The cell body of GFAP+ cells was segmented, and the total cell number and cell body occupied area were quantified. As astrocyte branches are thinner, more intercrossed, and denser than microglia, we first pronounced all the branches directly from images and then skeletonized them and quantified the total branch length in each view field. In the dentate gyrus (DG) of HIP (**Fig. 6a, Suppl. Table 4**), iTat had more astrocytes (**left panel, Fig. 6b**), smaller Cell Body Occupied Area (**middle panel, Fig. 6b**), and shorter Branch Length (**right panel, Fig. 6b**) than Wt in all groups, excepted for Branch Length in both males and females of 12m, Coc showed smaller Cell Body Occupied Area and shorter Branch Length than Sal in all groups, excepted for Branch Length in 6m males, and Coc showed even smaller Cell Body Occupied Area and shorter Branch Length than Sal in 6m iTat males, smaller Cell Body Occupied Area than Sal in 12m iTat females, and shorter Branch Lengths than Sal in 6m iTat females. There were much fewer astrocytes in CORT and CPU and the changes in these cells were not very evident (**data not shown**). These results showed that Tat activated astrocytes to proliferate, but with more naïve cells indicated by a smaller cell body and fewer branches, Coc only decreased astrocyte cell body and branch length, and Coc did not alter the number of astrocytes but further decreased astrocyte cell body and branch length. These findings suggest that Coc and Tat additively decreased the cell size and branches and diminished astrocyte activation (**Suppl. Table 5**).

**Figure 6.**
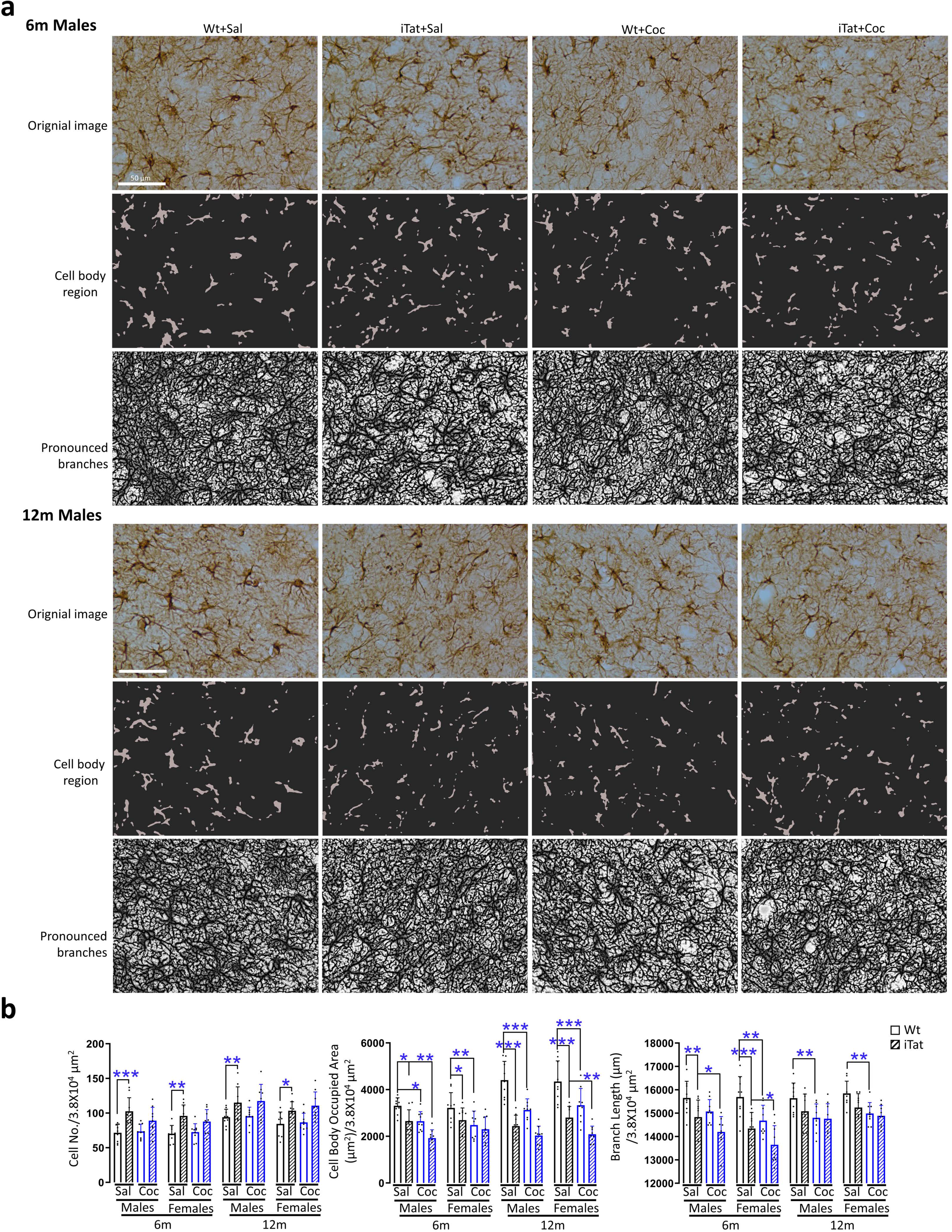
Effects of cocaine on astrocytes of HIP DG of iTat mice. Mouse brain sections were stained for GFAP, and the cell bodies of astrocytes in HIP DG were segmented and the branches were pronounced by Imaris and ilastik, respectively **(a**). The total number of astrocytes, the cell body occupied area, and the branch length were further calculated by Imaris or Cellprofiler in every view field (3.8 x 10^4^ μm^2^) (**b)**. Representative images were chosen from 6- and 12-months male mice groups. *P*<0.05 was considered significant and marked as *, and *p*<0.01 and *p*<0.001 were both considered highly significant and marked as ** and ***, respectively. Scale bars in **a**: 50 µm. Sal: saline, Coc: cocaine.

### Age and sex independently contributed to behavioral and pathological abnormalities by Tat and cocaine

Similar patterns of learning and memory and cellular response to Tat and cocaine were noted among all four groups of 2 ages x 2 sexes: 6m males, 6m females, 12m males, and 12m females (**Suppl. Table 1, 3 & 4**). To further ascertain if age and sex had different learning and memory and cellular response to Tat and cocaine, we performed statistical analysis for comparisons among these four groups (**Table 1**). In learning and memory of MWM, 12m females had longer Escape Latency and Cumulative Distance than 6m females, and 12m females showed longer Escape Latency and Cumulative Distance than 12m males, suggesting that the learning and memory of females are more vulnerable to ageing.

**Table 1:**
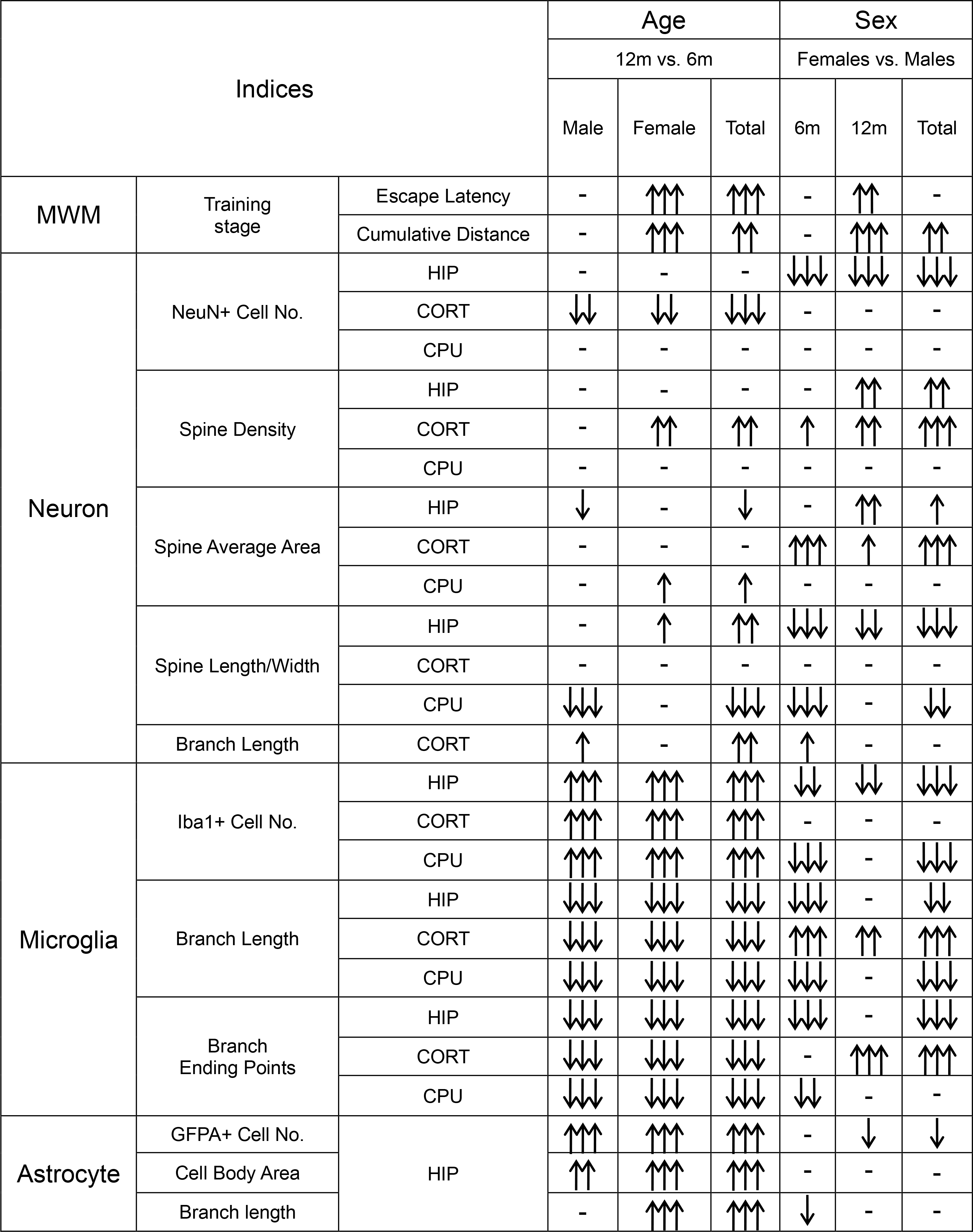
Summary of age and sex-related effects. One, two, and three arrows represented *p*<0.05, *p*<0.01, and *p*<0.001, respectively. Arrow down showed higher than the average value; Arrow down showed lower than the average value.

In cellular response, compared to 6m males/females, 12m males/females had fewer neurons in CORT but more microglia and astrocytes in HIP, CORT, and CPU, higher Spine Density in CORT, smaller Spine Average Area in HIP, larger Spine Average Area in CPU, higher Spine Length/Width ratio in HIP, lower Spine Length/Width ratio in CPU, longer branches in neurons of CORT and astrocytes of HIP astrocytes, shorter branches and fewer branch endpoints in microglia of HIP, CORT, and CPU, and larger Cell Body Area in astrocytes of HIP. Males and females only differed in the distribution of statistical significance between these two age groups. Females had the statistical significance in CORT Spine Density, CPU Spine Average Area, HIP Spine Length/Width ratio, and HIP Branch Length of astrocytes, whereas males had the statistical significance in HIP Spine Average Area, CPU Spine Length/Width ratio, and CORT Branch Length of neurons. These findings show that older mice lose functional neurons and gain more activated glia cells, while neurons grow more atypical spines and elongate branches to compensate the neuron loss, which likely contributes to the cognition decline of old mice detected in MWM.

In addition, compared to males, females had fewer neurons in HIP, higher Spine Density and Spine Average Area in HIP and CORT, lower Spine Length/Width ratio in HIP and CPU, longer branches in neurons of CORT, fewer microglia and shorter Branch Length and fewer Branch Endpoints in HIP and CPU, longer Branch Length and fewer Branch Endpoints in CORT, fewer astrocytes and shorter Branch Length in HIP. Both 6m and 12m mice mostly had the same response. 6m mice had statistical significance in Spine Length/Width ratio in CPU, microglia counts in HIP with Branch Length and Branch Endpoints, which were not noted in 12m mice. Conversely, 12m mice had the statistical significance in Spine Density and Spine Average Area in HIP, and microglia Branch Ending Points in CORT, which were not noted in 6m mice. There were some exceptions, such as neuron Branch Length in CORT, microglia Branch ending points in CPU, and astrocyte Branch Length in HIP did not show the significance in both males and females (Total), but only showed the significance in 6m mice that had opposite response compared to 12m mice. These findings show that females have fewer neurons, higher spine density, shorter but larger size of spines, longer neuronal branches, fewer activated glial cells, and microglia of diverse morphology in different brain regions, suggesting that glia cells may not be involved in the cognition vulnerability of females.

### Chronic exposure led to significant changes in genome-wide DNA methylation in the presence of Tat

Next, we performed the whole genome bisulfate sequencing (WGBS) with single base resolution and determined the impact of chronic cocaine and long-term Tat on genome-wide DNA methylation, focusing on methylated cytosine adjacent to a guanine (CpG) or other three nucleotides adenine, thymine and cytosine (CpH). We chose HIP hemispheres from the mice of eight 12m groups, selected three samples from each group, and extracted their DNA for 24 WGBS libraries, as HIP and 12m were the brain region and age that were most noted for the changes in learning and memory and cellular response by cocaine and Tat. Using the threshold of a minimum of six reads per a mC site, we identified a total of 39,312,240 CpG and 149,800,650 CpH sites. Initial analysis did not indicate any significant differences in the number and distribution of mC sites between males and females (Y chromosomes were excluded from analysis). This was consistent with our findings that both males and females showed generally similar trends in their learning and memory and cellular response to cocaine and Tat. Therefore, we decided to combine males and females to have four groups and focus on two factors cocaine and Tat (**Suppl. Fig. 9**).

We first compared the average methylation level of each CpG site among four groups. Cocaine showed lower mCpG than Sal in Wt mice, while iTat showed higher mCpG than Wt in Sal. In addition, there was still a significant interaction between Tat and cocaine, despite their trade-off effects (**Fig. 7a**). The mCpG sites showed similar distribution patterns in all 20 chromosomes among these four groups (**Fig. 7b**). To further identify differentially methylated regions (DMR), we framed six continuous mC sites as a calculated unit in preset regions and used the linear mixed model to determine the methylation value (M value) in three comparisons including Wt- Sal vs. iTat-Sal (Tat factor), Wt-Sal vs. Wt-Coc (Coc factor), and interaction (Tat x Coc factor). For mCpG-related DMR, 18371 hypermethylated DMR (Hyper-DMR) and 20195 hypomethylated DMR (Hypo-DMR) were identified for Tat (**left panel, Fig. 7c**), 22269 Hyper-DMR and 34308 Hypo-DMR were identified for Coc (**left panel, Fig. 7d**), and 14838 Hyper-DMR and 15800 Hypo-DMR were identified for Tat x Coc (**left panel, Fig. 7e**). Next, we traced these significant DMR back to chromosomes and located four lowest numbers of DMR at chromosome 16, 18, 19, and X and four highest numbers at chromosome 2, 5, 7, and 11 across these four groups (**right panels, Fig. 7c-d**). These results were consistent with the chromosome distribution of mouse active genes [81], indicating that the impact of Tat, Coc, or Tat x Coc on chromosomes is broad and non-selective. Lastly, we intersected the significant DMR of these three comparisons and found that 5232 DMR were overlapped in all of them, and 21292, 36744 and 11541 DMR were unique to Tat, Coc, and Tat x Coc, respectively (**Fig. 7f**).

**Figure 7.**
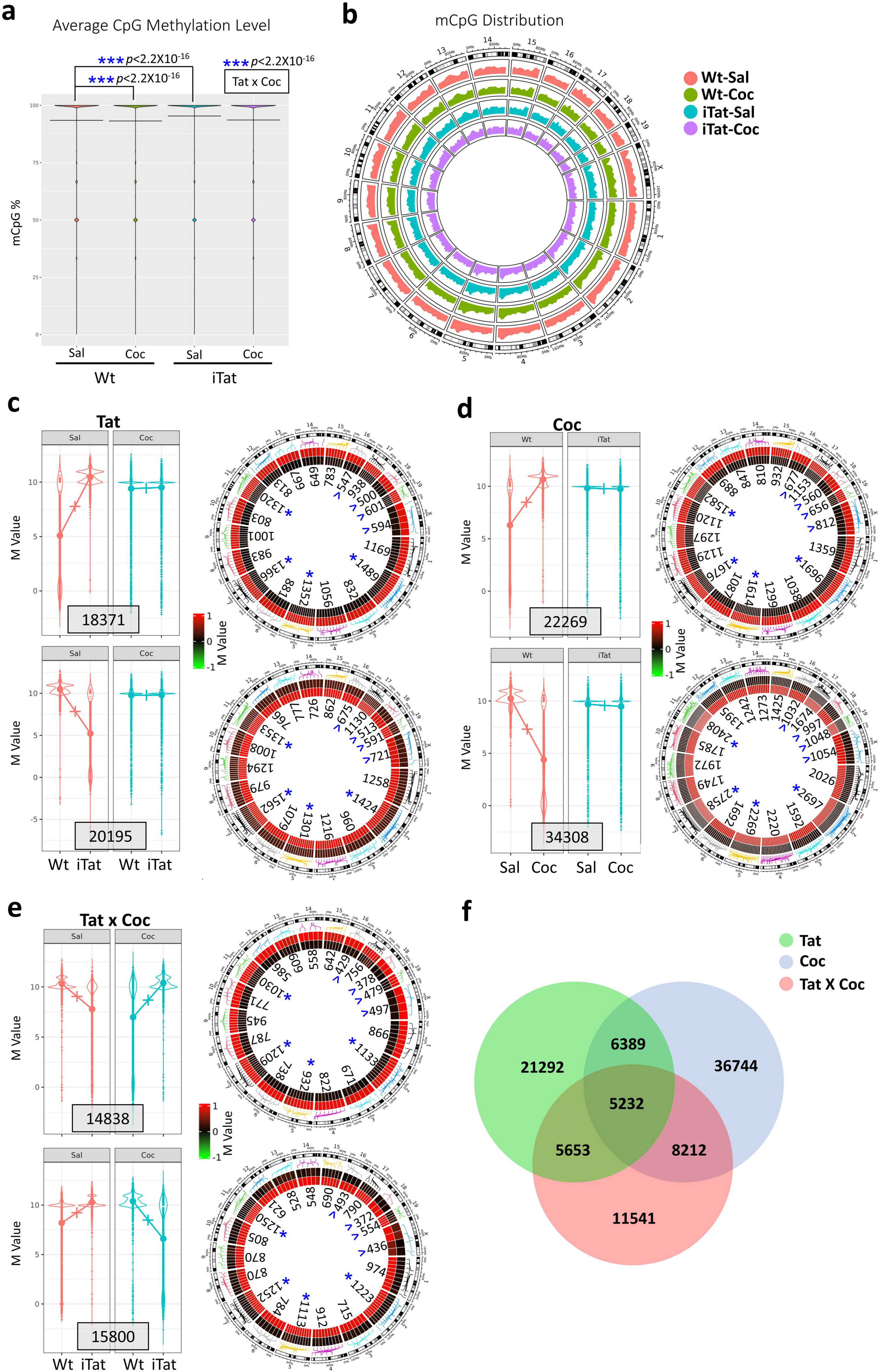
Effects of cocaine on genome-wide DNA methylation in the context of Tat expression. Genomic DNA was isolated from HIP of the mice for genome-wide DNA methylation analysis. **a**. The average CpG methylation level among these four groups; **b.** The CpG distribution among all mouse chromosomes. Red: Wt-Sal, Green: Wt-Coc, Blue: iTat-Sal, Pink: iTat-Coc. **c-e**. CpG sites-linked differentially methylated regions (DMR), expressed in M values (left panel) and their chromosomal location (right panel) under factor Tat (Wt-Sal vs. iTat- Sal, **c**), Coc (Wt-Sal vs. Wt-Coc, **d**), and Tat x Coc (interaction between Tat and Coc, **e**). *, Four chromosomes with the top ranked number of up and down DMR. **∨**, Four chromosomes with the bottom ranked total number of up and down DMR.

We also performed analysis of the average methylation level of each CpH site and chromosome distribution and obtained similar findings to CpG (**Suppl. Fig. 10a** & **b**). We then did the similar CpH-related DMR analysis. The total number of CpH-related Hyper-DMR and Hypo-DMR were lower than CpG-related Hyper-DMR and Hypo-DMR. 9234 Hyper-DMR and 3055 Hypo-DMR were identified for Tat (**left panel, Suppl. Fig. 10c**), 4427 Hyper-DMR and 7714 Hypo-DMR were identified for Coc (**left panel, Suppl. Fig. 10d**), and 1995 Hyper-DMR and 2762 Hypo-DMR were identified for Tat x Coc (**left panel, Suppl. Fig. 10e**). Chromosomes 16, 18, 19 and X had the lowest CpH-related DMR distribution for all factors which was same to CpG-DMRs (**right panels, Suppl. Fig. 10c-d**). However, four highest CpH-related DMR for these three factors showed different chromosomes, which were 1, 2, 4, 5, 6, and 7. There were 898 CpH- related DMR were overlapped in all three factors, and 8813, 8850, and 1860 CpH-related DMR were unique to Tat, Coc, and Tat x Coc, respectively (**Suppl. Fig. 10f**). We also investigated the distribution of CpG-DMR among promotors, exons, and introns relative to genomic distribution. However, no significant differences were found, even though the intron regions showed higher convergent tendency across three comparisons (**Suppl. Fig. 11a**). Similar findings were noted for CpH-related DMR for all three factors (**Suppl. Fig. 11b**). These results show that chronic cocaine and long-term Tat expression interactively led to significant changes of the genome-wide DNA methylation profiles, suggesting that epigenetic changes are likely involved in behavioral and cellular response to Tat and cocaine.

### Alterations of genome-wide gene expression by Tat and cocaine

To validate the DNA methylation changes by cocaine and Tat and to identify gene expression altered by these changes and linked to behavioral and cellular response to Tat and cocaine, we isolated RNA from the other corresponding HIP hemisphere of the same mice and performed bulky RNA sequencing (RNA-Seq). We used the same linear mixed model and identified differentially expressed genes (DEG) among these four groups and linked them with DMR (**Suppl. Fig. 9**). In CpG-related DMR, Tat altered 20,437 genes containing or proximal to Hyper-DMR and 22,583 for Hypo-DMR, of which 5,184 genes were overlapped; for Coc, 24,796 genes were containing or proximal to Hyper-DMR, 32,759 genes for Hypo-DMR, and 8,701 genes for both; for Tat x Coc, 17,950 genes were containing or proximal to Hyper-DMR, 18,518 genes for Hypo-DMR, and 3,959 genes for both (**Fig. 8a**). In addition, 513, 242, and 432 up-regulated DEG (Up-DEG) and 506, 479, and 186 down-regulated DEG (Down-DEG) were identified by Tat, Coc, and Tat X Coc, and 140, 79, and 127 Up-DEG were linked to CpG-related Hypo-DMR and 143, 143, and 52 Down-DEG were linked to CpG-related Hyper-DMR under Tat, Coc, and Tat X Coc factor, respectively.

**Figure 8.**
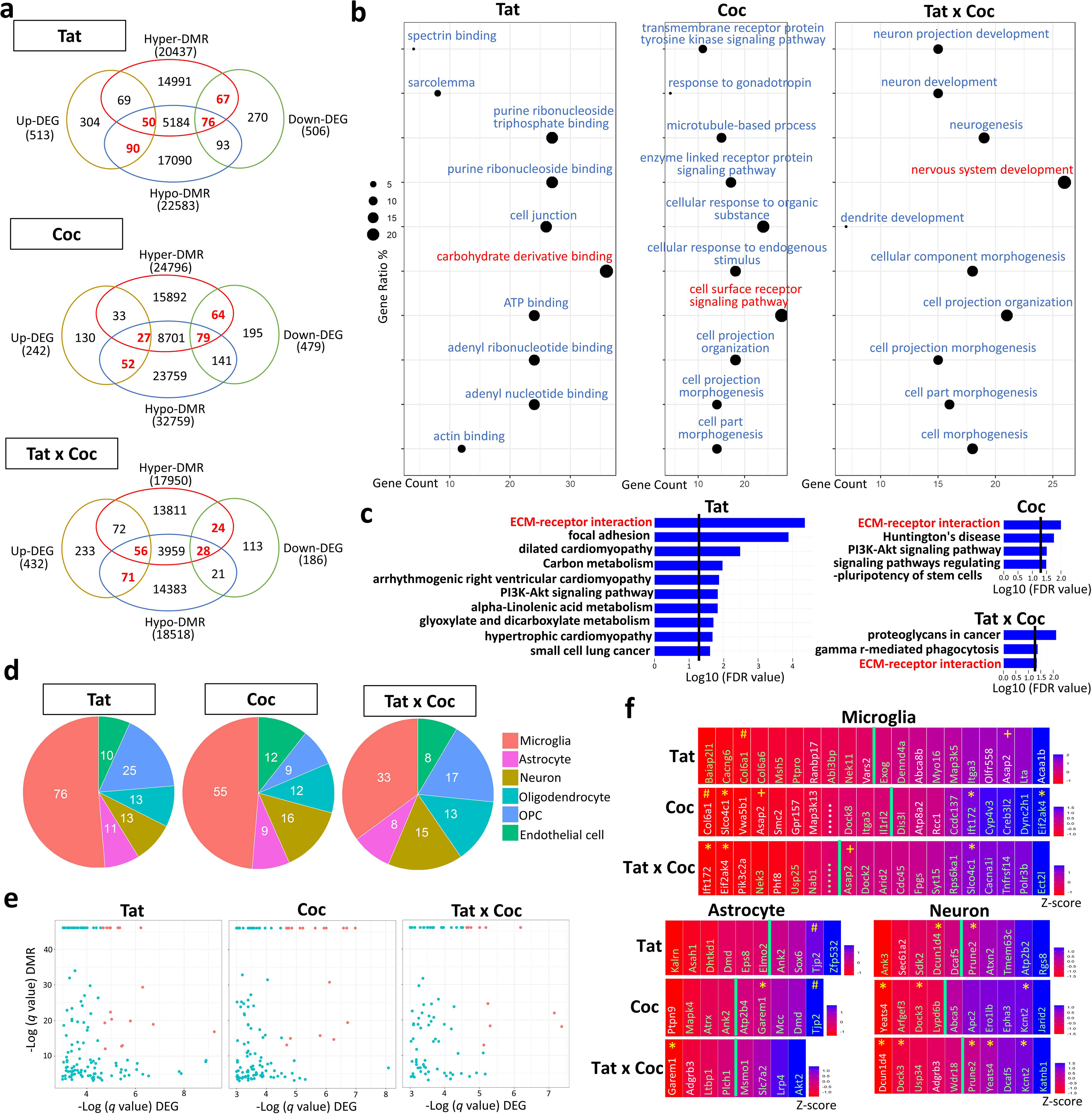
The relationship between CpG DNA methylation and genes expression by each of the factors Tat, Coc, and Tat x Coc. RNA was isolated from HIP of the mice and used for RNA-Seq analysis. Differentially expressed genes (DEG, Up or Down) were identified to be proximal to DMR (Hyper or Hypo) by each of the factors Tat, Coc, and Tat x Coc (**a**). Up- DEG/Hypo-DMR-linked genes and Down-DEG/Hyper-DMR-linked genes (marked in red in **a**) were chosen to run GO (**b**) and KEGG enrichment analysis (**c**). The GO cluster with the highest gene ratio and gene count was highlighted in red (**b**). The extracellular matrix (ECM)-receptor interaction pathway was shared among these three factors (marked in red, **c**). DEG in each factor were further segregated to different brain cell types: microglia, astrocyte, neuron, oligodendrocyte, oligodendrocyte progenitor cell (OPC), and endothelial cell (**d**). The most significantly impacted genes that had *q*<1e-5 DMR and *q*<0.01 DEG were marked in red in volcano map (**e**). Top 20 DEG with *q*<0.05 in microglia and top 10 DEG with *q*<0.05 in astrocytes and neurons were shown in heatmap (**f**), in which overlapped top genes were marked in white. White ellipsis: 9330182L06Rik; Green ellipsis: 4933407I05Rik. *, genes present in both Tat x Coc and Tat or Coc; #, genes present in both Tat and Coc; +, genes present in Tat, Coc, and Tat x Coc.

Next, we performed GO and KEGG enrichment analysis on these overlapped genes to determine whether there were any specific functions or pathways involved. Tat, Coc, and Tat x Coc had different top 10 functional clusters (*p* < 0.05) from GO functional analysis (**Fig. 8b**). The cluster with the largest gene ratio and gene counts for Tat was *Carbohydrate derivative binding*, for Coc was *cell surface receptor signaling pathway*, and for Tat x Coc was *nervous system development*. Several morphology-related clusters were overlapped between Coc and Tat x Coc, such as *cell projection morphogenesis*, *cell part morphogenesis*, and *cell projection organization*, suggesting cocaine plays a pivotal role in inducing pathological changes (**middle and right panels, Fig. 8b**). More neuron functional clusters occurred in Tat x Coc, including *neuron projection development*, *neuron development*, *neurogenesis,* and *dendrite development* (**right panel, Fig. 8b**), which provides molecular evidence to support our behavioral findings that chronic cocaine use led to more severe cognition decline in iTat mice. Different to GO functional clusters, KEGG pathway analysis revealed more convergent effects among these three factors, as *extracellular matrix (ECM)-receptor interaction* pathway was the found in all of them (**Fig. 8c**). ECM is deeply involved in synapse formation and plasticity [82]. These findings suggest that Tat, Coc, and Tat x Coc affect neuronal function commonly through *ECM-receptor interaction* pathway.

To determine the roles of different cell types in response to Tat and cocaine, we employed machine learning to build up the database of different brain cell types and then used this database to tag the above overlapped genes into Neuron, Microglia, Astrocyte, oligodendrocyte, oligodendrocyte progenitor cell (OPT), and endothelial cell. Specifically, we took the top 6000 ranked cell type-enriched mouse genes [74], mapped them with the RNA-Seq transcriptome database from purified brain cells [75], employed Support Vector Machine [76] to build a learning model with these mapped genes, and applied the trained model to predict the cell type for the unmapped genes. The largest number of genes were in microglia for all three factors (**Fig. 8d**), suggesting that microglia was the most affected cell type by Tat, Coc, and Tat x Coc. All other 5 cell types appeared to be somewhat equally affected among these three factors.

To identify the most promising gene targets for the morphological changes of microglia, astrocyte, and neuron and mouse behavior, we used volcano map to sort out genes with *q*< 1e-5 DMR and *q*<0.01 DEG, marked by red (**Fig. 8e**) and heatmap to document these 20 top ranked genes in microglia and top ranked 10 genes, if there were, in astrocyte and neuron with *q*<0.05 DEG (**Fig. 8f**). The genes marked in red in volcano map were highlighted in white in heatmap, and these genes had the highest significance in both DNA methylation and gene expression by Tat, Coc, and Tat x Coc. Down-regulation of *Ranbp17* and *Vars2* and up-regulation of *Abca8b, Olfr558, Asap2, and Acaa1b* in microglia and *Sec61a2f* in neuron were highly involved in Tat-induced behavioral and pathological changes; for Coc, down-regulation of *Col6a1, Slco4c1, Vwa5b1, Asap2, Smc2, Gpr157, Map3k13,* and *9330182L06Rik* in microglia, *Ptpn9* in astrocytes, and *Yeats4* in neuron and up-regulation of *Atp8a2, Rcc1* in microglia; and for Tat x Coc, down- regulation of gene *Ift172*, *Eif2ak4*, *Pik3c2a*, *Phf8* in microglia, *Garem1* and *Adgrb3* in astrocytes, and *Dcun1d4* and *Adgrb3* in neuron.

Other genes, which were not highlighted in heatmap, could also play an important role in behavioral and cellular changes by these three factors, as they all reached significant standards of DMR (*q*<0.05) (**Fig. 8f**). Of particular note was that several genes were present in at least two of these three factors Tat, Coc and Tat x Coc. In microglia, *Ift172* and *Eif2ak4* were down-regulated in Tat x Coc, but showed up-regulated in Coc. *Slco4c1* showed ug-regulated in Tat x Coc, but down-regulated in Coc; and, *Asap2* showed up-regulated in Tat and Tat x Coc but down- regulated in Coc. In astrocytes, *Garem1* was down-regulated Tat x Coc, but up-regulated in Coc. In neuron, *Dcun1d4* and *Prune2* were up-regulated and down-regulated, respectively, in both Tat x Coc and Tat, but to a different extent; Similarly, *Dock3* and *Kcnt2* were up-regulated and down-regulated, respectively, in both Tat x Coc and Coc; and *Yeats4* was up-regulated in Tat x Coc but down-regulated in Coc. In addition, genes like *Col6a1* in microglia and *Tjp2* in astrocytes were either up-regulated or down-regulated in both Tat and Coc factors, indicating that they may have additive effect between Tat and cocaine.

For CpH-related DMR, there were 9375 genes containing or proximal to Hyper-DMR and 3670 for Hypo-DMR by Tat, and 631 genes were overlapped for both; for Coc, 5379 genes were containing or proximal to Hyper-DMR, 8293 genes for Hypo-DMR, and 822 genes for both; for Tat x Coc, 2381 genes were containing or proximal to Hyper-DMR, 3188 genes for Hypo-DMR, and 187 genes for both (**Suppl. Fig. 12a**). In addition, there were 31, 35, and 14 Up-DEG linked to Hypo-DMR and 62, 50, and 19 Down-DEG linked to Hyper-DMRs by Tat, Coc, and Tat x Coc factors, respectively. Microglia was also most affected by all these factors, while other 5 cell types showed similar effects among these three factors (**Suppl. Fig. 12b**). However, only a small number of genes were screened out for all three factors, which did not allow sufficient statistic power for further GO and KEGG enrichment analysis and volcano map and heatmap selection.

## DISCUSSION

### Interactive effects between Tat and cocaine

Cocaine exerted its impact on the behaviors of iTat mice in an interactive manner and on glial and neuronal response by Tat in both interactive and additive manner (**Suppl. Table 4**). Specifically, MWM indices with significant changes were all interactive effect (**Suppl. Table 1**). For morphological changes in neurons, SYP and PSD-95 expression levels had either interactive or additive effect, differing among the groups, and the spine morphology changes all showed interactive effect (**Suppl. Table 2**); For morphological changes in microglia, all showed interactive effect, while for morphological changes in astrocyte, all showed additive effect (**Suppl. Table 3**). In agreement with our studies, other studies show interactive or additive effect between cocaine and Tat, including cholesterol homeostasis of glia and neurons [57, 58], excitability of pyramidal neurons in medial prefrontal cortex [55], neuronal differentiation [62], brain energy metabolism [59, 60], and addiction [83]. However, few of these studies have clearly clarified the differences between interactive and additive effect. Although the both interactive and additive effect could lead to worse outcomes, interactive effect is biologically more significant and often leads to more unpredictable outcomes.

In terms of neurobehaviors and neuropathology, Tat slightly impaired learning and memory (**Fig. 1b** & **c**, **Suppl. Fig. 1**, **Suppl. Table 1**) and altered spine morphology (**Fig. 2a** & **b**, **Suppl. Fig. 3**, **4a** & **b, Suppl. Table 2**). In contrast, cocaine showed better learning and memory, although there was no statistical significance when compared to Sal. Cocaine was also associated with higher spine density, average area, and length/width ratio. However, Tat and cocaine together showed significantly worsened learning and memory and induced dramatic spine dysgenesis (swelling). Similarly, in HIP of 6m males and 12m females, Tat barely affected SYP expression, while cocaine increased SYP expression; However, Tat and cocaine together drastically elevated SYP expression (**Fig. 4, Suppl. Table 2**). In microglia morphology, Tat significantly increased branch length and the number of branch endpoints, while cocaine had similar but to much less extent; However, Tat and cocaine together led to remarkably decreased branch length and the branch endpoints (**Fig. 5**, **Suppl. Fig. 7** & **8, Suppl. Table 3**). Furthermore, 14838 Hyper- and 15800 Hypo-CpG DMR linked to 56 down-regulated and 127 up-regulated DEG (**Fig. 7e** & **Fig. 8a**), and 1995 Hyper- and 2762 Hypo-CpH DMR linked to 19 down-regulated and 14 up- regulated DEG (**Suppl. Fig. 10e** & **Suppl. Fig. 12a**) were identified to be involved in the interactions between Tat and cocaine. All these findings strongly support the interactive nature between Tat and cocaine for the impact of cocaine in the HAND population.

Besides the interactions between Tat and cocaine, chronic cocaine use enhances spatial learning and memory [84, 85]. Cocaine leads to higher spine density [86–88] and alteration of spine morphology [89, 90]. Chronic cocaine use also enhances long-term potentiation in different brain regions [91–93]. Thus, spine morphology is reasonably linked to the learning and memory. In this study, consistent with other studies [94–97], we showed that changes of synaptic markers SYP and PSD-95 expression by cocaine varied among brain regions, age and sex (**Fig. 4**, **Suppl. Fig. 5**, **6a & b, Suppl. Table 2**), indicating that effects of cocaine are context-dependent. This possibility is supported by other behavioral findings in the study that cocaine unexpectedly and specifically showed somewhat anxiolytic effect on 6m females by EPM and OPT (**Suppl. Fig. 2a** & **b, Suppl. Table 5**) and anti-depressive effect to 12m males by TST and FST (**Suppl. Fig. 2d & e, Suppl. Table 5**). Also, as demonstrated in other studies [98, 99], we showed cocaine activated microglia in 6m mice with increased branch length and ending points and decreased astrocyte cell body regions and branch lengths (**Fig. 5** & **6**, **Suppl. Fig. 7** & **8, Suppl. Table 3**).

We and others have shown that Tat impairs spatial memory and locomotor activity [47, 64] and causes depressive status [100, 101], neuronal cell loss [102, 103], shortened neuron dendrite branches [104], lower spine density [105], and glia cells activation [64, 103], which we also showed in this study (**Suppl. Table 1-5**). However, we provided the first direct and *in vivo* evidences that Tat caused neuronal loss in HIP, CORT and CPU, shortened neuron dendrite branches in CORT, lower spine density in HIP and CPU, microglia activation along with increased cell number and branches in HIP, CORT, and CPU, and astrocyte activation with increased cell number and decreased cell body and branch length in HIP. To our knowledge, this is the first study to delineate the involvement of age and sex factors in changes of behavioral and cellular response to Tat and cocaine (**Table 1, Suppl. Table 1, 3-5**). Age and sex (female) mostly showed additively effects. However, there was an exception, that is that significant depressive status was noted in 6m males by FST and somewhat depressive status in 12m females by both TST and FST (**Suppl. Fig. 2d** & **e, Suppl. Table 5**), indicating that age and sex function by different mechanisms. Another finding of interest was that all iTat mice showed higher anxious threshold (anxiolytic characteristic) in EPM and OPT even with cocaine (**Suppl. Fig. 2a** & **b, Suppl. Table 5**). It is possible that the locus coeruleus reactivity of the anxiety system [106] could be dysregulated by Tat. This possibility merits further investigation.

### Glia involvement in synaptic dysgenesis

Neuronal synapse remodeling occurs throughout lifetime, which is a necessary and sufficient procedure for memory encoding and storage [107, 108]. Microglia and astrocytes proactively participate in synapse formation (establishment) and pruning (maintenance), which comprise the most important part of synaptic plasticity. Usually, astrocytes ensheath synapses, maintain neurotransmitter hemostasis and deliver the regulation signals to microglia; once microglia receive signals from astrocytes or other cell types, it will start phagocytosis process to induce engulfment of synapses [109–112]. As such, astrocyte is an important functional compartment for synapses, while microglia, a highly motile surveillant type of cells serve as a synapse-pruning player. We found that chronic cocaine use sharply diminished branch lengths of microglia and astrocytes along with synapse swelling in iTat mice (**Fig. 2**, **5**, **6**, **Suppl. Fig. 3**, **4**, **7**, **8, Supp. Table 3** & **4**). A plausible explanation is that chronic cocaine use limits astrocyte-neuron contact and damages microglia pruning process, which in turn leads to unrestrained synapse formation resulting synapse swelling or dysgenesis to have enough capacity to mediate impaired learning and memory process, which our findings supported (**Fig. 1b** & **c**, **Suppl. Fig. 1**, **Suppl. Table 1**). In addition, ECM that constitutes around 20% brain volume has been proposed to be a potential mediator for astrocytes-microglia-synapse interaction involving in synapse remodeling [82, 113]. Our KEGG pathway analysis, which showed *ECM-receptor interaction* pathway was significantly involved in all three factors Tat-, cocaine-, and Tat/cocaine-induced molecular changes (**Fig. 8c**), offers additional evidence to further support this possibility. Our GO functional analysis revealed that more neuron functional and morphological clusters were present under Tat x cocaine factor (**Fig. 8b**), which could be the molecular basis for synaptic dysgenesis.

Of note is that although astrocytes play an important role during this process, the significant proportion of microglia-related genes with significant changes in their expression levels were identified in our bioinformatic analysis (**Fig. 8d**). Lastly, the morphology changes of microglia showed the interactive effect between Tat and cocaine, which was different from additive effects in astrocytes (**Suppl. Table 5**). All these evidences support the notion that microglia is the major player during synaptic dysgenesis by Tat and cocaine.

### Roles of genome-wide DNA Methylation by Tat and cocaine

Most studies about cocaine-associated DNA methylation changes focus on nucleus accumbens and prefrontal cortex of the brain and show different levels of genome-wide DNA methylation by different approaches (review [114]). We have previously shown that Tat decreases genome-wide DNA methylation in CORT and cerebellum by ELISA [47]. In this study, we determined the genome-wide DNA methylation in HIP using the single base resolution WGBS. Tat or Tat and cocaine decreased the CpG and CpH percentage, cocaine had no significant changes in the CpG percentage and slight increases in the CpH percentage (**Suppl. Fig. 13**), suggesting that Tat alters the mC sites and cocaine does not. However, Tat increased the average methylation levels of each CpG and CpH site, and cocaine showed opposite effects (**Fig. 7a**, **Suppl. Fig. 10a**). Taken together, Tat reduced the numbers of CpG and CpH sites, but increased their average methylation level. In contrary, cocaine did not affect the numbers of CpG and CpH sites but decreased their average methylation level. These findings suggest that the changes at individual CpG and CpH site may be more valuable than the changes at the genome-wide level. In addition, we showed that the distribution of these mCpG and mCpH sites and their linked DMR by Tat, cocaine or Tat and cocaine did NOT show significant convergent effect (**Fig. 7b-e**, **Suppl. Fig. 10b-e** & **11**), indicating that Tat and cocaine hijack upstream genes/pathways that control DNA methylation. DNMT3B, which we identified to be responsive to Tat [47], could be a strong candidate.

### Tat as an accelerating factor of biological aging

In the study, we monitored body weights of all mice and found that iTat mice weighed significantly lower than Wt mice in both 6m and 12m groups (**Suppl. Fig. 14a**), which extended our previous observation that there were weight differences between Wt and iTat mice of one month [26]. We also found that females weighed significantly lower than males in 6m iTat and Wt mice as well as 12m Wt mice, but not in 12m iTat mice, suggesting that iTat mice could experience andropause and menopause earlier than Wt mice or their sex hormone could decline earlier than Wt mice. In addition, we noticed that 12 m iTat mice had much higher mortality than 12m Wt mice during cocaine injection (**Suppl. Table 6**), although there was no statistical significance due to the small group number. Nonetheless, these results indicate that aging iTat mice are more vulnerable than Wt mice to stress, such as injection, cocaine or the declined adaptability of the cardiovascular system to stress earlier than Wt mice. To ascertain this Tat- ageing connection [47], we aligned HIP mC sites, which were generated from WGBS in 12m mice, with 732 documented age-related mC markers [115]. We identified 172 matched mC sites and found that the methylation level (M value) of 32 mC sites in iTat mice were significantly higher from Wt mice (**Suppl. Fig. 15, Suppl. Table 7**). These results further demonstrated that iTat mice had higher biological age and support that Tat accelerates biological aging.

## Supporting information

All Supplementary materials

## ACKNOWLEDGEMENTS

This work was supported in part by grants R01DA043162 and R01NS094108 (JJH) and P20GM103449 from the National Institutes of Health.

