## Supplementary material for "Cocaine exacerbates neurological impairments and neuropathologies in the iTat model of HIV-associated neurocognitive disorder through genome-wide alterations of DNA methylation and gene expression": All Supplementary materials

**Supplementary Table 1. Summary of changes in MWM by Tat and cocaine**

| Stage | index | Tat | | | | Coc | | | | Tat x Coc | | | |
|---|---|---|---|---|---|---|---|---|---|---|---|---|---|
| | | iTat+Sal vs Wt+Sal or iTat-Total vs Wt-Total | | | | Wt+Coc vs Wt+Sal or Coc-Total vs Sal-Total | | | | iTat+Coc vs iTat+Sal or Wt+Coc | | | |
| | | 6m-M | 12m-M | 6m-F | 12m-F | 6m-M | 12m-M | 6m-F | 12m-F | 6m-M | 12m-M | 6m-F | 12m-F |
| Training stage | Escape Latency | - | - | - | - | - | - | - | - | ↑↑ | ↑ | ↑↑ | - |
| | Cumulative Distance | - | - | ↑ | - | - | - | - | - | ↑↑ | - | ↑↑ | - |
| 1st probe test | Latency to Platform | - | - | - | - | - | - | - | - | - | - | - | - |
| | Platform Entries | ↓ | - | - | - | - | - | - | - | - | - | - | - |
| | Time at Target Quadrant | - | - | - | - | - | - | - | - | ↓ | | - | - |
| | Distance to Target Quadrant | - | - | - | - | - | - | - | - | ↓ | | - | - |
| | Time at Platform | ↓ | - | - | - | - | - | - | - | - | - | - | - |
| | Distance to Platform | - | - | - | - | - | - | - | - | - | - | - | - |
| 2nd probe test | Latency to Platform | - | ↑ | - | - | - | - | - | - | - | - | - | - |
| | Platform Entries | - | - | - | - | - | - | - | - | - | ↓↓ | - | - |
| | Time at Target Quadrant | - | - | - | - | - | - | - | - | - | ↓↓ | ↓↓ | - |
| | Distance to Target Quadrant | - | - | - | - | - | - | - | - | ↓ | | - | - |
| | Time at Platform | - | - | - | - | - | - | - | - | - | - | - | - |
| | Distance to Platform | - | - | - | - | - | - | - | - | - | - | - | - |

One, two and three arrows represented *p*<0.05, *p*<0.01, and *p*<0.001, respectively. Arrow up showed higher than the average value; Arrow down showed lower than the average value.

**Supplementary Table 2. Summary of changes of other behaviors by Tat and cocaine**

| Test | Index | Tat | | | | Coc | | | | Tat x Coc | | | |
|---|---|---|---|---|---|---|---|---|---|---|---|---|---|
| | | iTat+Sal vs Wt+Sal or iTat-Total vs Wt-Total | | | | Wt+Coc vs Wt+Sal or Coc-Total vs Sal-Total | | | | iTat+Coc vs iTat+Sal or Wt+Coc | | | |
| | | 6m-M | 6m-F | 12m-M | 12m-F | 6m-M | 6m-F | 12m-M | 12m-F | 6m-M | 6m-F | 12m-M | 12m-F |
| EPM | Open Arm Entries | ↑ | ↑ | - | - | - | ↑↑ | - | - | ↑ | - | - | ↑ x |
| | Open Arm Time | - | - | ↑↑↑ | - | - | - | - | - | - | - | - | - |
| | Open Arm Distance | - | - | ↑↑ | - | - | ↑ | - | - | ↑ | - | - | ↑ x |
| OPT | Total Distance | - | - | - | ↓ | - | - | - | - | - | ↑ x | - | - |
| | Maximum Speed | - | ↓ | - | ↓↓↓ | - | - | - | - | - | - | - | ↑ x |
| | Central Distance | - | - | ↑ | - | - | - | ↑ | - | - | ↑↑ x | - | - |
| | Central Entries | - | - | ↑↑ | - | - | ↑ | - | - | - | ↑↑ x | - | - |
| RT | Latency to Fall (30 rpm) | | ↓ | / | / | - | - | / | / | - | - | / | / |
| | Latency to Fall (15 rpm) | / | / | - | - | / | / | - | - | / | / | - | ↑ x |
| FST | Immobile Time | ↑ | - | - | - | - | - | ↓ | - | - | - | - | - |
| TST | Immobile Time | - | - | - | - | - | - | - | - | - | - | - | ↓ x |

One, two and three arrows represented $p<0.05$, $p<0.01$, and $p<0.001$, respectively. Arrow up showed higher than the average value; Arrow down showed lower than the average value. Green x: no interactive effect and only additive effect between Tat and cocaine. Black x: no interactive or additive effect between Tat and cocaine, and iTat+Coc group maintains the same pattern to Wt+Coc group.

**Supplementary Table 3. Summary of changes of the number of neurons and neuronal morphology by Tat and cocaine**

| Cell type | Index | Brain region | Tat | | | | Coc | | | | Tat x Coc | | | |
|---|---|---|---|---|---|---|---|---|---|---|---|---|---|---|
| | | | iTat+Sal vs Wt+Sal or iTat-Total vs Wt-Total | | | | Wt+Coc vs Wt+Sal or Coc-Total vs Sal-Total | | | | iTat+Coc vs iTat+Sal or Wt+Coc | | | |
| | | | 6m-M | 6m-F | 12m-M | 12m-F | 6m-M | 6m-F | 12m-M | 12m-F | 6m-M | 6m-F | 12m-M | 12m-F |
| Neuron | NeuN+ Cell No. | HIP | ↓↓↓ | ↓↓↓ | ↓↓↓ | ↓↓↓ | - | - | - | - | - | - | - | - |
| | | CORT | ↓↓↓ | - | ↓↓ | - | - | - | - | - | - | - | - | - |
| | | CPU | ↓↓↓ | ↓↓↓ | ↓↓ | ↓↓ | - | - | - | - | - | - | - | - |
| | Spine Density | HIP | - | - | - | - | ↑↑ | ↑↑↑ | ↑ | ↑↑↑ | - | ↑↑↑ | - | - |
| | | CORT | - | - | - | - | ↑ | ↑↑↑ | ↑ | ↑ | - | - | ↑↑↑ | ↑ |
| | | CPU | ↓↓ | - | - | - | - | ↑ | ↑ | ↑ | ↑ | - | - | - |
| | Spine Average Area | HIP | - | ↓ | - | - | ↑ | - | - | - | - | - | ↑↑↑ | ↑ |
| | | CORT | - | - | - | - | - | - | ↑ | - | ↑↑ | ↑↑↑ | ↑↑ | ↑↑↑ |
| | | CPU | - | - | - | - | ↑↑ | - | ↑↑↑ | ↑ | ↑↑↑ | - | ↑↑↑ | - |
| | Spine Length/Width | HIP | - | ↓↓↓ | - | - | ↑ | - | - | - | - | - | - | - |
| | | CORT | - | - | - | ↓↓↓ | - | ↑↑ | ↑↑ | - | - | - | - | - |
| | | CPU | ↑↑↑ | - | - | - | ↑↑↑ | - | - | - | ↓↓↓ | - | ↓↓ | - |
| | Branch Length | CORT | ↓↓ | - | - | - | - | - | - | - | - | - | - | - |
| | SYP Pre-synaptic marker | HIP | - | - | - | - | ↑↑↑ | - | ↑↑↑ | ↑↑ | ↑↑↑ | ↑ x | ↑ x | ↑↑↑ |
| | | CORT | - | - | - | ↑ | - | - | ↑ | ↑ | ↑ x | - | ↑ x | ↑ x |
| | | CPU | - | - | - | - | - | - | - | - | ↑ x | - | ↑ x | ↑ x |
| | PSD-95 Pre-synaptic marker | HIP | - | - | - | - | ↓↓↓ | ↓↓ | ↓↓ | ↓↓ | ↑ | ↑↑↑ | ↑ | ↑ |
| | | CORT | - | - | - | ↓ | - | - | - | - | ↑↑ | ↑↑ | - | - |
| | | CPU | - | - | - | - | - | - | ↓↓ | - | - | ↑↑ x | - | - |

One, two and three arrows represented $p<0.05$, $p<0.01$, and $p<0.001$, respectively. Arrow up showed higher than the average value; Arrow down showed lower than the average value. Green x: no interactive effect and only additive effect between Tat and cocaine.

**Supplementary Table 4. Summary of changes of the number of astrocytes and their morphology by Tat and cocaine**

| Cell type | Index | Brain region | Tat | | | | Coc | | | | Tat x Coc | | | |
|---|---|---|---|---|---|---|---|---|---|---|---|---|---|---|
| | | | iTat+Sal vs Wt+Sal or iTat-Total vs Wt-Total | | | | Wt+Coc vs Wt+Sal or Coc-Total vs Sal-Total | | | | iTat+Coc vs iTat+Sal or Wt+Coc | | | |
| | | | 6m-M | 6m-F | 12m-M | 12m-F | 6m-M | 6m-F | 12m-M | 12m-F | 6m-M | 6m-F | 12m-M | 12m-F |
| Microglia | Iba1+ Cell No. | HIP | ↑↑↑ | ↑ | ↑ | - | - | - | - | - | - | - | - | - |
| | | CORT | ↑↑↑ | ↑↑↑ | ↑↑↑ | ↑ | - | - | - | - | - | - | - | - |
| | | CPU | ↑↑↑ | ↑↑ | ↑↑↑ | ↑ | - | - | - | - | - | - | - | - |
| | Branch Length | HIP | ↑↑↑ | ↑↑ | ↑ | ↑ | ↑↑↑ | ↑↑ | - | - | ↓↓↓ | ↓↓↓ | ↓↓↓ | - |
| | | CORT | ↑ | ↑↑ | ↑↑↑ | - | - | - | - | - | ↓↓↓ | ↓↓↓ | ↓↓↓ | - |
| | | CPU | ↑↑↑ | ↑↑↑ | - | ↑ | ↑↑ | ↑↑↑ | - | - | - | ↓↓↓ | ↓↓ | - |
| | Branch Ending Points | HIP | ↑↑↑ | ↑↑ | ↑ | ↑↑ | ↑↑ | ↑↑ | - | - | ↓↓↓ | ↓↓↓ | ↓↓↓ | - |
| | | CORT | ↑↑↑ | ↑↑ | ↑↑↑ | - | ↑ | - | - | - | ↓↓↓ | ↓↓↓ | ↓↓↓ | - |
| | | CPU | ↑↑↑ | ↑↑↑ | ↑ | ↑ | ↑ | ↑↑↑ | - | - | - | ↓↓ | ↓↓↓ | - |
| Astrocyte | GFPA+ Cell No. | HIP | ↑↑↑ | ↑↑ | ↑↑ | ↑ | - | - | - | - | - | - | - | - |
| | Cell Body Area | | ↓ | ↓ | ↓↓↓ | ↓↓↓ | ↓ | ↓↓ | ↓↓↓ | ↓↓↓ | ↓↓ x | - | - | ↓↓ x |
| | Branch Length | | ↓↓ | ↓↓↓ | - | - | - | ↓↓ | ↓↓ | ↓↓ | ↓ x | ↓ x | - | - |

One, two and three arrows represented $p<0.05$, $p<0.01$, and $p<0.001$, respectively. Arrow up showed higher than the average value; Arrow down showed lower than the average value. Green x: no interactive effect and only additive effect between Tat and cocaine.

**Supplementary Table 5. Interactive and additive effect between Tat and cocaine**

| Tat X Coc | MWM | | Neuron | | | | | | | Microglia | | | Astrocytes | | |
|---|---|---|---|---|---|---|---|---|---|---|---|---|---|---|---|
| | Training stage | Probe Test | Cell No. | Spine | | | Branch | Synapse | | Cell No. | Branch | | Cell No. | Cell Body | Branch |
| | | | NeuN+ | Density | Area | Length/width | Length | Pre (SYP) | Post (PSD) | Iba+ | Length | Ending point | GFAP+ | Area | length |
| *Interactive | Yes | Yes | No | Yes | | | No | Yes | Yes | No | Yes | | No | No | No |
| *Additive | No | No | No | No | | | No | Yes | Yes | No | No | | No | Yes | Yes |

***,** Interactive effect was defined by ANOVA tests or Linear mix model analysis. If there was no interactive effect and combined effect of these two factors was more than each of the factors alone, it was defined as additive effect.

**Supplementary Table 6: Mouse mortality during injection**

| Groups | | Died mice No. | | Total No. | Mortality |
|---|---|---|---|---|---|
| Strain | Sex | Sal | Coc | | |
| Wt | Male | - | 1 | 23 | 2.3% |
| | Female | - | - | 21 | |
| iTat | Male | 1 | 3 | 28 | 10.3% |
| | Female | - | 2 | 30 | |

**Supplementary Table 7. ID of mC sites with significant differences**

| ID | q value |
|---|---|
| 4_139581246 | 2.54E-19 |
| 19_45741556 | 2.44E-07 |
| 4_139581142 | 1.69E-05 |
| 2_146959272 | 0.000102 |
| 8_108714475 | 0.000109 |
| 2_30798754 | 0.000177 |
| 10_81643336 | 0.000321 |
| 11_102606283 | 0.000387 |
| 2_146959254 | 0.000389 |
| 4_99653882 | 0.000389 |
| 8_105271015 | 0.000453 |
| 11_57832738 | 0.000472 |
| 3_55782956 | 0.000472 |
| 7_133776161 | 0.000529 |
| 8_108886958 | 0.00066 |
| 19_45741606 | 0.00066 |
| 16_33382347 | 0.000688 |
| 15_98780554 | 0.000688 |
| 8_105271004 | 0.000863 |
| 12_69196914 | 0.000997 |
| 12_69196918 | 0.001613 |
| 11_57832763 | 0.001672 |
| 12_111442906 | 0.001773 |
| 4_139581155 | 0.001778 |
| 12_111028824 | 0.001778 |

| | |
|---|---|
| 6_30959010 | 0.002384 |
| 6_30959001 | 0.00333 |
| 8_125669949 | 0.005361 |
| 6_30959032 | 0.013046 |
| 15_98780491 | 0.013243 |
| 8_125670015 | 0.017949 |
| 6_88190216 | 0.020864 |

### SUPPLEMENTARY FIGURE LEGENDS

**Supplementary Figure 1**. **Extended MWZ data from Fig. 1b.** Cumulative Distance to platform during the MWZ training stage was also recorded (**a**). The second probe test was performed to measure the long-term memory of mice 7 days after the first probe test, and all indices used were the same as **Fig. 1b**. $p<0.05$ was considered significant and marked as *, # or $ for comparisons among different groups; $p<0.01$ and $p<0.001$ were both considered highly significant and marked as ** and ***, respectively. Sal: saline, Coc: cocaine.

**Supplementary Figure. 2**: **Effects of cocaine on other behaviors of iTat mice. a.** EPM was performed on day 25 to measure the anxious status of mice and expressed by three indices: Open Arm Entries, Open Arm Time and Open Arm Distance. **b**. OPT was performed on day 26 to measure the locomotor activity and expressed by three indices: Total Distance, Maximum Speed, and Central Distance and Central entries. **c**. RT was performed on day 27 for the accelerated mode and day 28 for the fixed mode. After the accelerated mode screening, 30 rpm was set up as sensitive mode for 6-month groups (6m) and 15 rpm for 12-month groups (12m), and the Latency to fall at those speeds was recorded. **d & e.** TST and FST were performed on day 29 and 30, respectively, to measure the depressive status of mice. The immobile time during the entire experimental process (6 min) was chosen in TST (d), while only the last four min of immobile time was used as the index for FST (**e**). $p<0.05$ was considered significant and marked as *, and $p<0.01$ and $p<0.001$ were both considered highly significant and marked as ** and ***, respectively. Sal: saline, Coc: cocaine.

**Supplementary Figure 3**. **Effects of cocaine on dendritic spine morphology and the number of neurons in CORT of iTat mice. a** & **b**. Mouse brain sections were stained in the Golgi-Cox solution. z stack images of dendrites and axons of neurons in CORT were taken, projected in the same layer by Fiji, and segmented out all branches by ilastik (**a**). The branch length in each individual cell was calculated by Cellprofiler (**b**). **c** & **d**. Mouse brain sections were stained for NeuN and counter stained in DAPI (**c**) and the number of NeuN-positive neurons in CORT were quantified and expressed as a fraction of the number of the DAPI-positive cells in CORT (**d**). Representative images were chosen from 6- and 12-month males (**b** & **c**). $p<0.05$ was considered significant and marked as *, and $p<0.01$ and $p<0.001$ were both considered highly significant and marked as ** and ***, respectively. Scale bars: 10 μm for **a**, 100 μm for **c**. Sal: saline, Coc: cocaine.

**Supplementary Figure 4**. **Effects of cocaine on dendritic spine morphology and the number of neurons in striatum (CPU) of iTat mice. a** & **b**. Mouse brain sections were stained in the Golgi-Cox solution. z stack images of dendrites and axons of neurons in CPU were taken, projected in the same layer by Fiji, and segmented out all branches by ilastik (**a**). The branch length in each individual cell was calculated by Cellprofiler (**b**). **c** & **d**. Mouse brain sections were stained for NeuN and counter stained in DAPI (**c**) and the number of NeuN-positive neurons in CPU were quantified and expressed as a fraction of the number of the DAPI-positive cells in CPU (**d**). Representative images were chosen from 6- and 12-month males (**b** & **c**). $p<0.05$ was considered significant and marked as *, and $p<0.01$ and $p<0.001$ were both considered highly significant and marked as ** and ***, respectively. Scale bars: 10 μm for **a**, 100 μm for **c**. Sal: saline, Coc: cocaine.

**Supplementary Figure 5**. **Effects of cocaine on synaptophysin expression in cortex (CORT) of iTat mice.** The CORT regions were dissected from the mouse brain and processed for lysates. SYP and PSD-95 expression in the lysates were determined by Western blotting (**a**), quantified by Fiji, normalized to the loading control β-actin, and calculated using the Wt+Sal as a reference, which was set at 1 (**b**). $p<0.05$ was considered significant and marked as *, and $p<0.01$ and $p<0.001$ were both considered highly significant and marked as ** and ***, respectively. Sal: saline, Coc: cocaine.

**Supplementary Figure 6**. **Effects of cocaine synaptophysin expression in caudate putnam (CPU) of iTat mice.** The CPU region were dissected from the mouse brain and processed for lysates. SYP and PSD-95 expression in the lysates were determined by Western blotting (**a**), quantified by Fiji, normalized to the loading control β-actin, and calculated using the Wt+Sal as a reference, which was set at 1 (**b**). $p<0.05$ was considered significant and marked as *, and $p<0.01$ and $p<0.001$ were both considered highly significant and marked as ** and ***, respectively. Sal: saline, Coc: cocaine.

**Supplementary Figure 7**. **Effects of cocaine on microglia of CORT of iTat mice.** Mouse brain sections were stained for Iba-1, the cell bodies of microglia were segmented and the branches were skeletonized by Cellprofiler (**a**). The total number of microglia in every view field (1.4 x $10^5$ $\mu m^2$), and the branch length of microglia and the number of the ending points in each microglia were further calculated by Cellprofiler (**b**). Representative images were chosen from parietal cortex in 6- and 12-month males. $P<0.05$ was considered significant and marked as *, and $p<0.01$ and $p<0.001$ were both considered highly significant and marked as ** and ***, respectively. Scale bars in **a**: 50 μm. Sal: saline, Coc: cocaine.

**Supplementary Figure. 8**: **Effects of cocaine on microglia of CPU of iTat mice.** Mouse brain sections were stained for Iba-1, the cell bodies of microglia were segmented and the branches were skeletonized by Cellprofiler (**a**). The total number of microglia in every view field (1.4 x $10^5$ $\mu m^2$), and the branch length of microglia and the number of the ending points in each microglia were further calculated by Cellprofiler (**b**). Representative images were chosen from 6- and 12-month males. $P<0.05$ was considered significant and marked as *, and $p<0.01$ and $p<0.001$ were both considered highly significant and marked as ** and ***, respectively. Scale bars in **a**: 50 μm. Sal: saline, Coc: cocaine.

**Supplementary Figure 9**. Bioinformatic analysis pipeline. 24 HIP hemispheres of 12-month male and female mice (three samples/group, total eight groups) was chosen for whole genome bisulfate sequencing (WGBS). After initial analysis, the sex factor was removed as there were no significant differences between male and female mice, and eight groups were then merged to four group including Wt-Sal, iTat-Sal, Wt-Coc, and iTat-Coc. A linear mix model was used to analyze the differential methylated regions (DMR). In parallel, the corresponding HIP hemispheres from the same mice were processed for bulk RNA sequencing (RNA-Seq). The same linear mix model was used to identify the Differential Gene Expression (DEG). Then, DMR were linked to proximal DEG to determine their potential target regulatory genes.

**Supplementary Figure 10**. **Effects of cocaine on genome-wide DNA methylation in the context of Tat expression.** Genomic DNA was isolated from HIP of the mice for genome-wide DNA methylation analysis. **a**. The average CpH methylation level among these four groups; **b.** The CpH distribution among all mouse chromosomes**.** Red: Wt-Sal, Green: Wt-Coc, Blue: iTat-Sal, Pink: iTat-Coc. **c-e**. CpH sites-linked differentially methylated regions (DMR), expressed in M values (left panel) and their chromosomal location (right panel) under factor Tat (Wt-Sal vs. iTat-Sal, **c**), Coc (Wt-Sal vs. Wt-Coc, **d**), and Tat x Coc (interaction between Tat and Coc, **e**). *, Four chromosomes with the top ranked number of hyper- and hypo-DMR. ∨, Four chromosomes with the bottom ranked total number of hyper- and hypo-DMR.

**Supplementary Figure 11. Distribution of CpG and CpH-related DMR among promoters, exons, and introns**. The fold enrichment score is the percentage of DMR in different genomic locations including Promotor, Exon, and Intron were divided by the percentage of reference genomic location type, and dashed line at y = 1 indicates the genomic values. **a**. CpG-related DMR. **b**. CpH-related DMR. The *p* values for all of them were higher than 0.05.

**Supplementary Figure 12**. **The relationship between CpH DNA methylation and genes expression by each of the factors Tat, Coc, and Tat x Coc.** RNA was isolated from HIP of the mice and used for RNA-Seq analysis. Differentially expressed genes (DEG, Up or Down) were identified to be proximal to DMR (Hyper or Hypo) by each of the factors Tat, Coc, and Tat x Coc (**a**). DEG in each factor were further segregated to different brain cell types: microglia, astrocyte, neuron, oligodendrocyte, oligodendrocyte progenitor cell (OPC), and endothelial cell (**b**).

**Supplementary Figure 13**. **Comparisons between the number of CpG and the number of CpH among these four experimental groups.** A total of 39,312,240 CpG and 149,800,650 CpH sites were identified after deduplication across all samples. The percentage of CpG (**a**) and CpH (**b**) in each group was calculated by the number of CpG or CpH sites in each group divided by the number of CpG or CpH sites in all four groups.

**Supplementary Figure 14**. **Changes of body weight of Wt and iTat mice.** All mice were weighted at the beginning of cocaine injection (**a**) and monitored for weight changes during the 14-day cocaine injection (**b**). $P<0.05$ was considered significant and marked as *, and $p<0.01$ and $p<0.001$ were considered highly significant and marked as ** and ***, respectively.

**Supplementary Figure 15. Ageing-related mC sites in HIP of iTat mice.** HIP mC sites were aligned with 732 documented age-related mC markers, 172 ageing-related mC sites were identified, and the differences of their methylation level between Wt and iTat groups were analyzed (**a**), and 32 mC sites with significances were shown by heatmap (**b**).

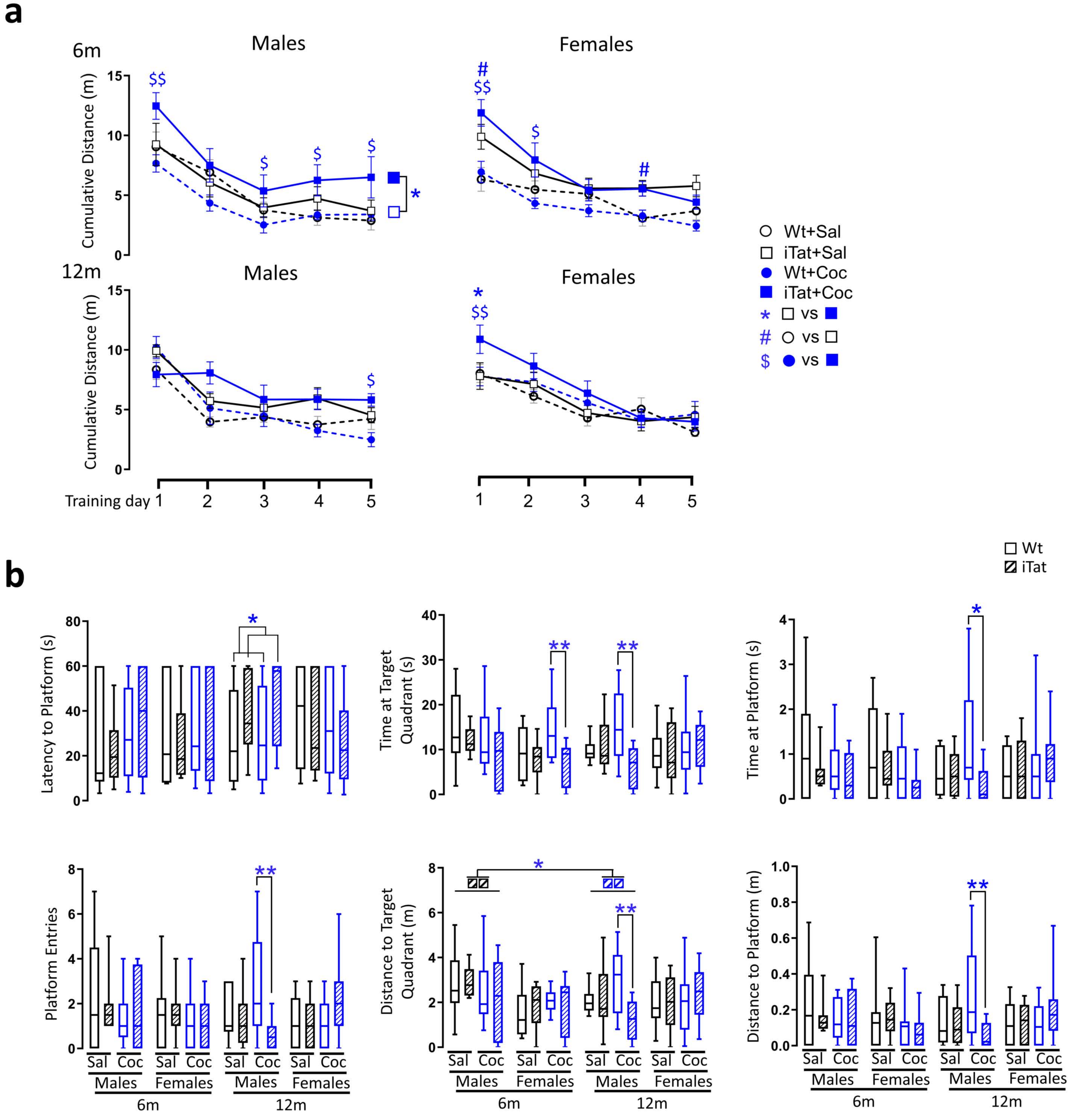
a
6m
Males
Females
12m
Males
Females
Cumulative Distance (m)
Training day
Wt+Sal
iTat+Sal
Wt+Coc
iTat+Coc
* vs
# vs
$ vs
b
Wt
iTat
Latency to Platform (s)
Time at Target Quadrant (s)
Time at Platform (s)
Platform Entries
Distance to Target Quadrant (m)
Distance to Platform (m)
Sal
Coc
Males
Females
6m
12m

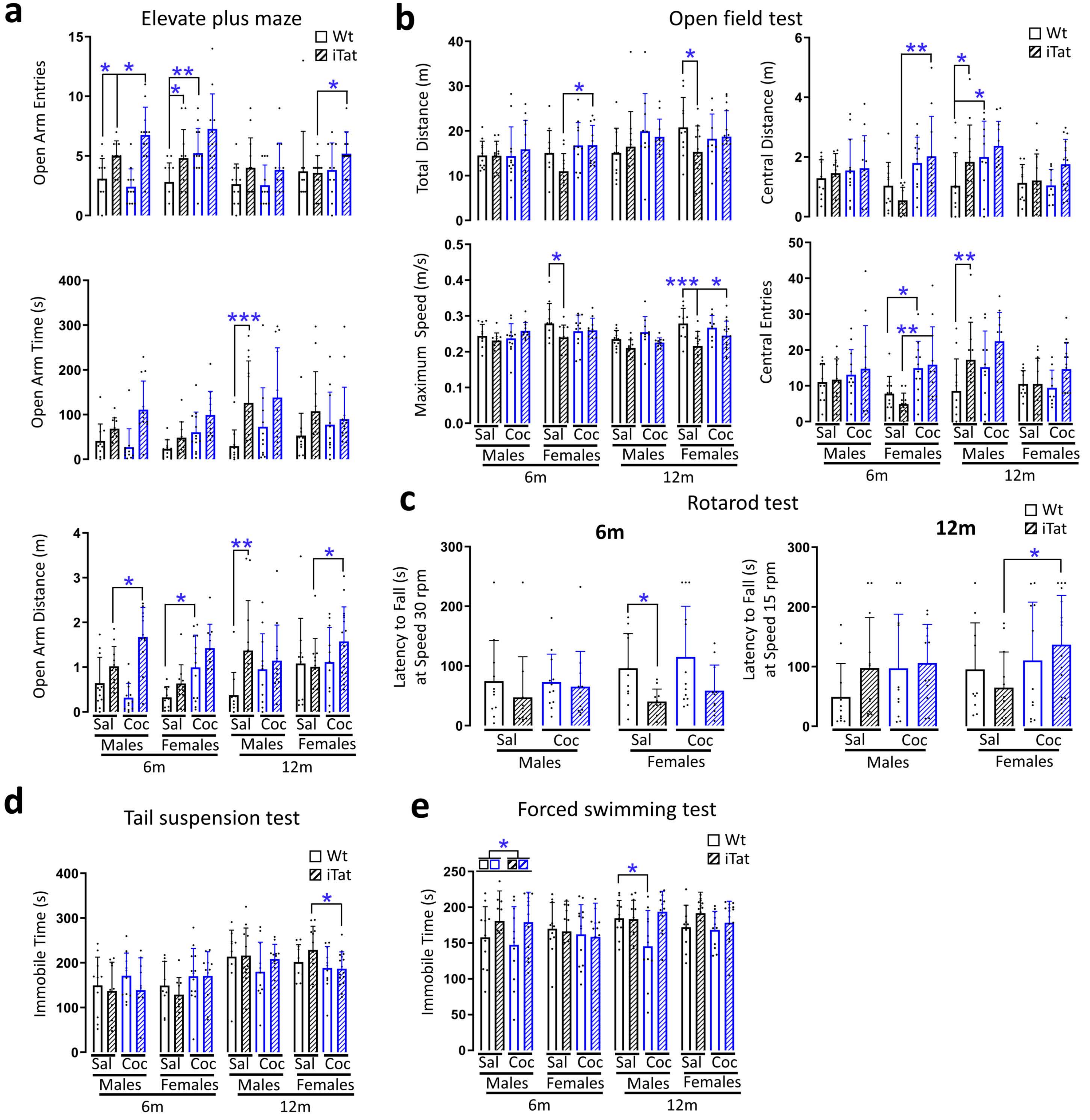

a
Elevate plus maze
Wt
iTat
Open Arm Entries
Open Arm Time (s)
Open Arm Distance (m)
Sal Coc Sal Coc Sal Coc Sal Coc
Males Females Males Females
6m 12m
b
Open field test
Total Distance (m)
Central Distance (m)
Maximum Speed (m/s)
Central Entries
c
Rotarod test
6m
12m
Latency to Fall (s) at Speed 30 rpm
Latency to Fall (s) at Speed 15 rpm
d
Tail suspension test
Immobile Time (s)
e
Forced swimming test
Immobile Time (s)

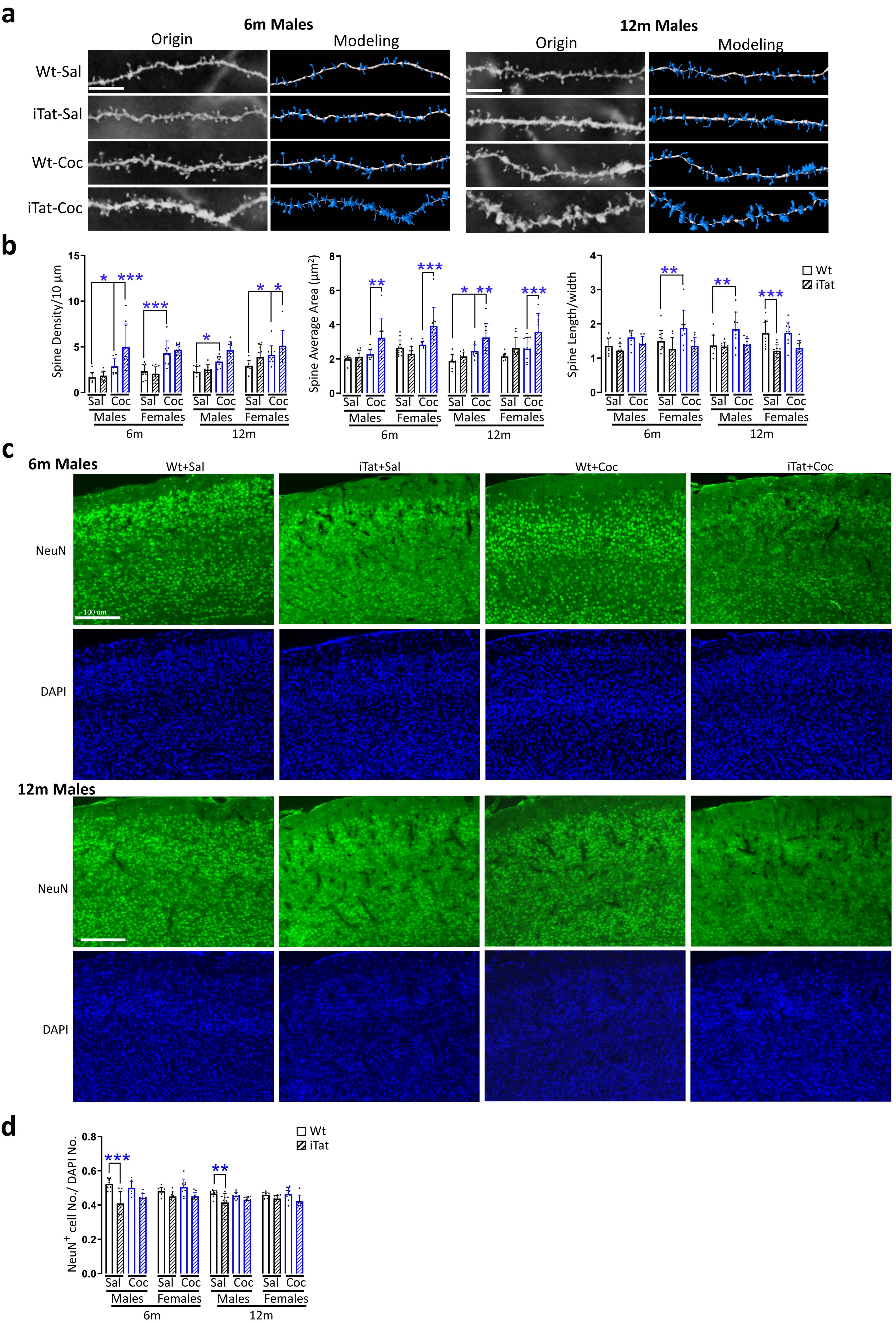

a
6m Males
Origin
Modeling
12m Males
Origin
Modeling
Wt-Sal
iTat-Sal
Wt-Coc
iTat-Coc
b
Spine Density/10 μm
Spine Average Area (μm2)
Spine Length/width
Wt
iTat
Sal Coc Sal Coc Sal Coc Sal Coc
Males Females Males Females
6m
12m
c
6m Males
Wt+Sal
iTat+Sal
Wt+Coc
iTat+Coc
NeuN
DAPI
100 um
12m Males
NeuN
DAPI
d
NeuN+ cell No./ DAPI No.
Wt
iTat
Sal Coc Sal Coc Sal Coc Sal Coc
Males Females Males Females
6m
12m

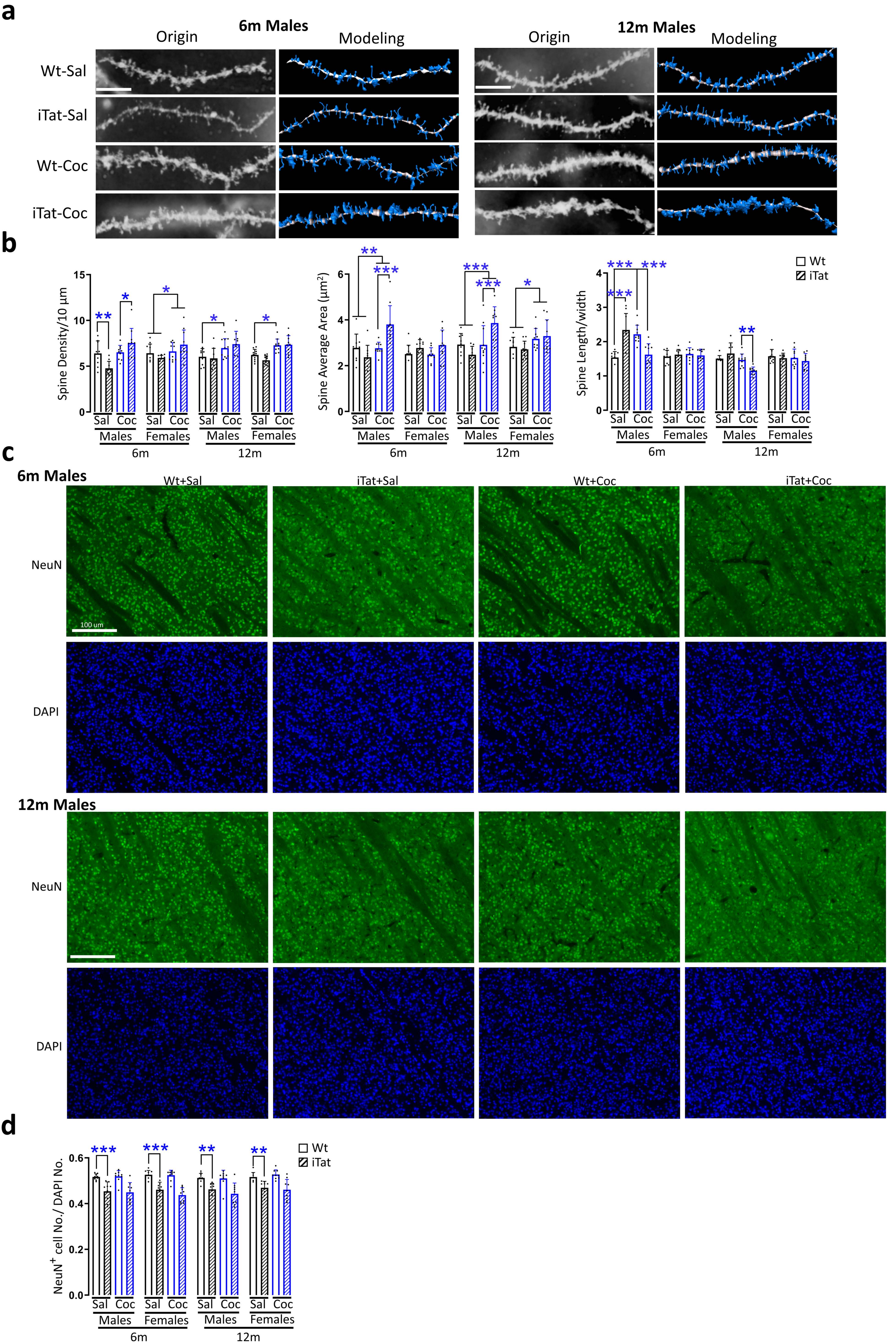
a
6m Males
Origin
Modeling
12m Males
Origin
Modeling
Wt-Sal
iTat-Sal
Wt-Coc
iTat-Coc
b
Spine Density/10 µm
Spine Average Area (µm²)
Spine Length/width
Wt
iTat
Sal Coc Sal Coc Sal Coc Sal Coc
Males Females Males Females
6m 12m
c
6m Males
Wt+Sal
iTat+Sal
Wt+Coc
iTat+Coc
NeuN
DAPI
100 um
12m Males
NeuN
DAPI
d
NeuN+ cell No./ DAPI No.
Wt
iTat
Sal Coc Sal Coc Sal Coc Sal Coc
Males Females Males Females
6m 12m

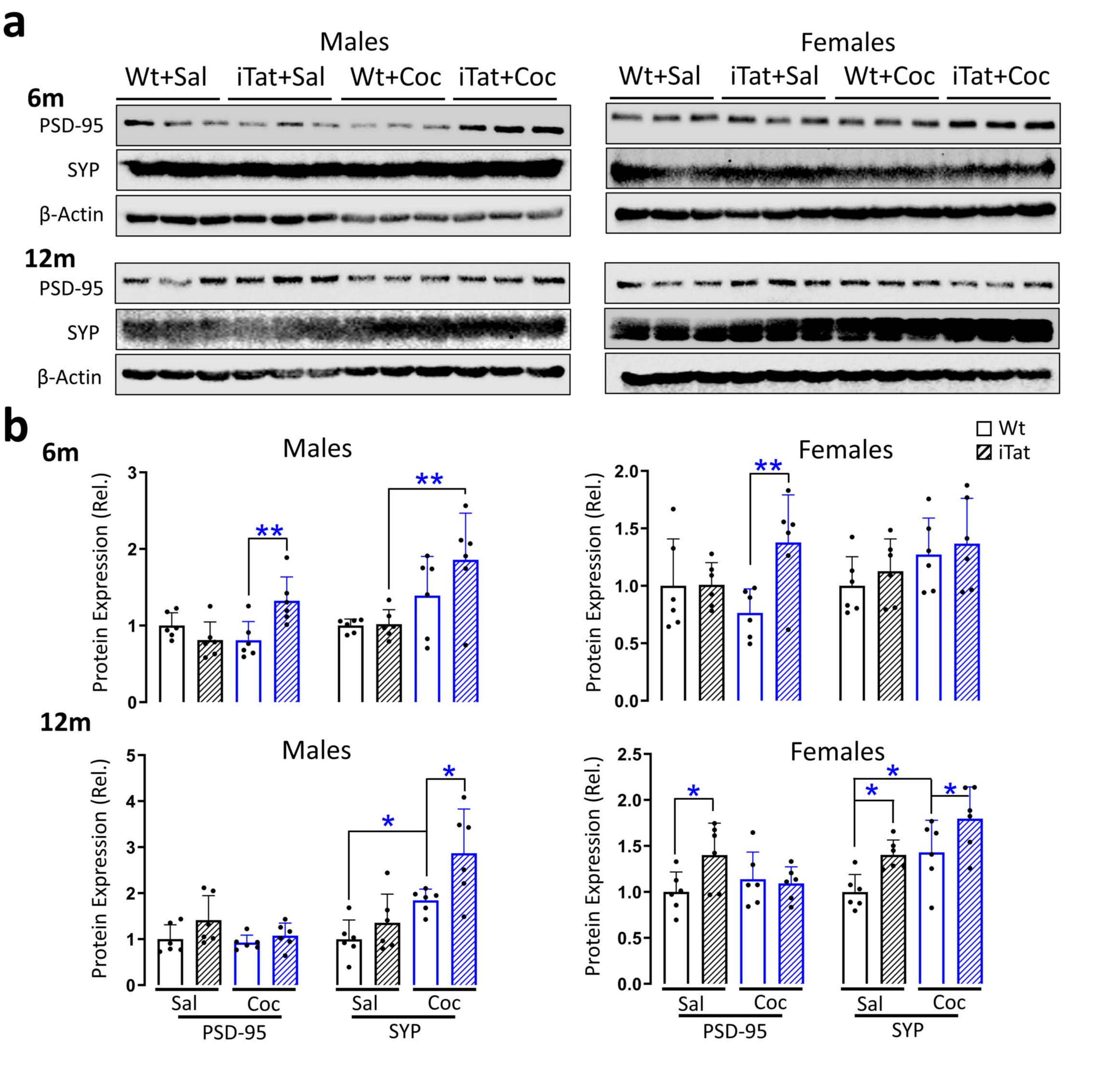
a
Males
Females
Wt+Sal
iTat+Sal
Wt+Coc
iTat+Coc
6m
PSD-95
SYP
β-Actin
12m
b
Wt
iTat
Protein Expression (Rel.)
Sal
Coc
PSD-95
SYP

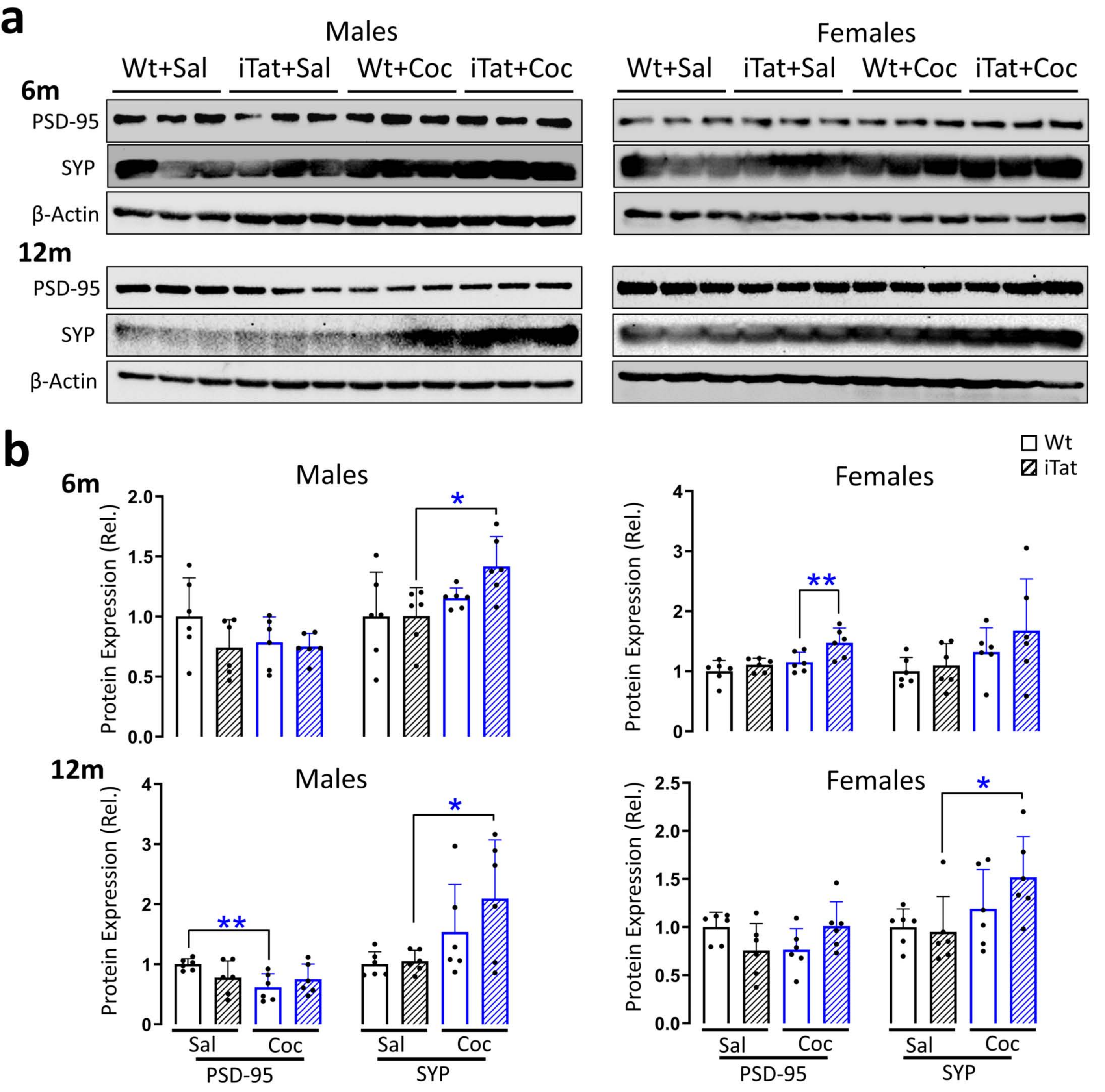

a
Males
Females
Wt+Sal
iTat+Sal
Wt+Coc
iTat+Coc
6m
12m
PSD-95
SYP
β-Actin
b
Wt
iTat
Protein Expression (Rel.)
Sal
Coc
PSD-95
SYP

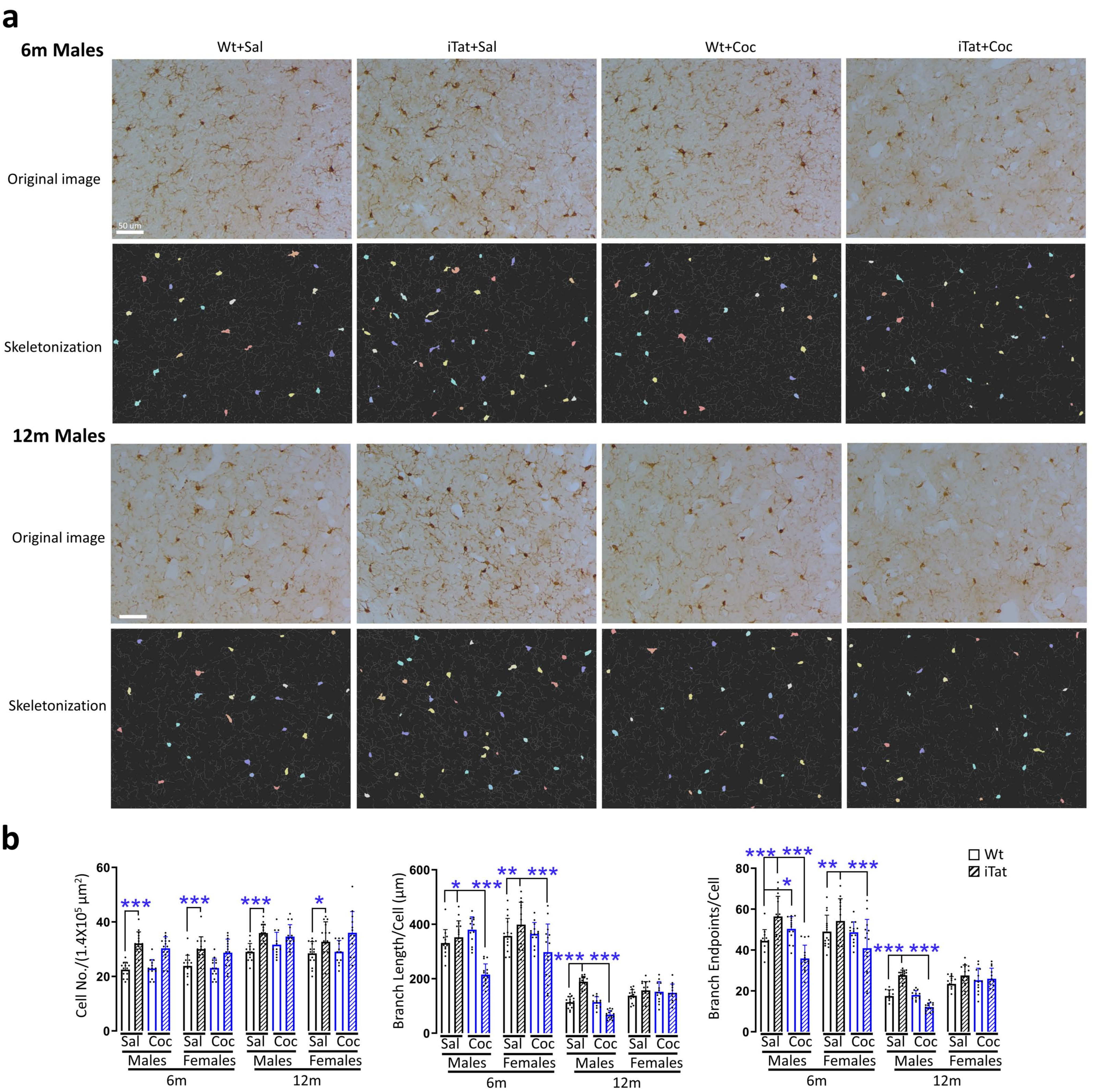
a
6m Males
Wt+Sal
iTat+Sal
Wt+Coc
iTat+Coc
Original image
50 um
Skeletonization
12m Males
Original image
Skeletonization
b
Cell No./(1.4X10^5 μm^2)
Branch Length/Cell (μm)
Branch Endpoints/Cell
Wt
iTat
Sal
Coc
Males
Females
6m
12m

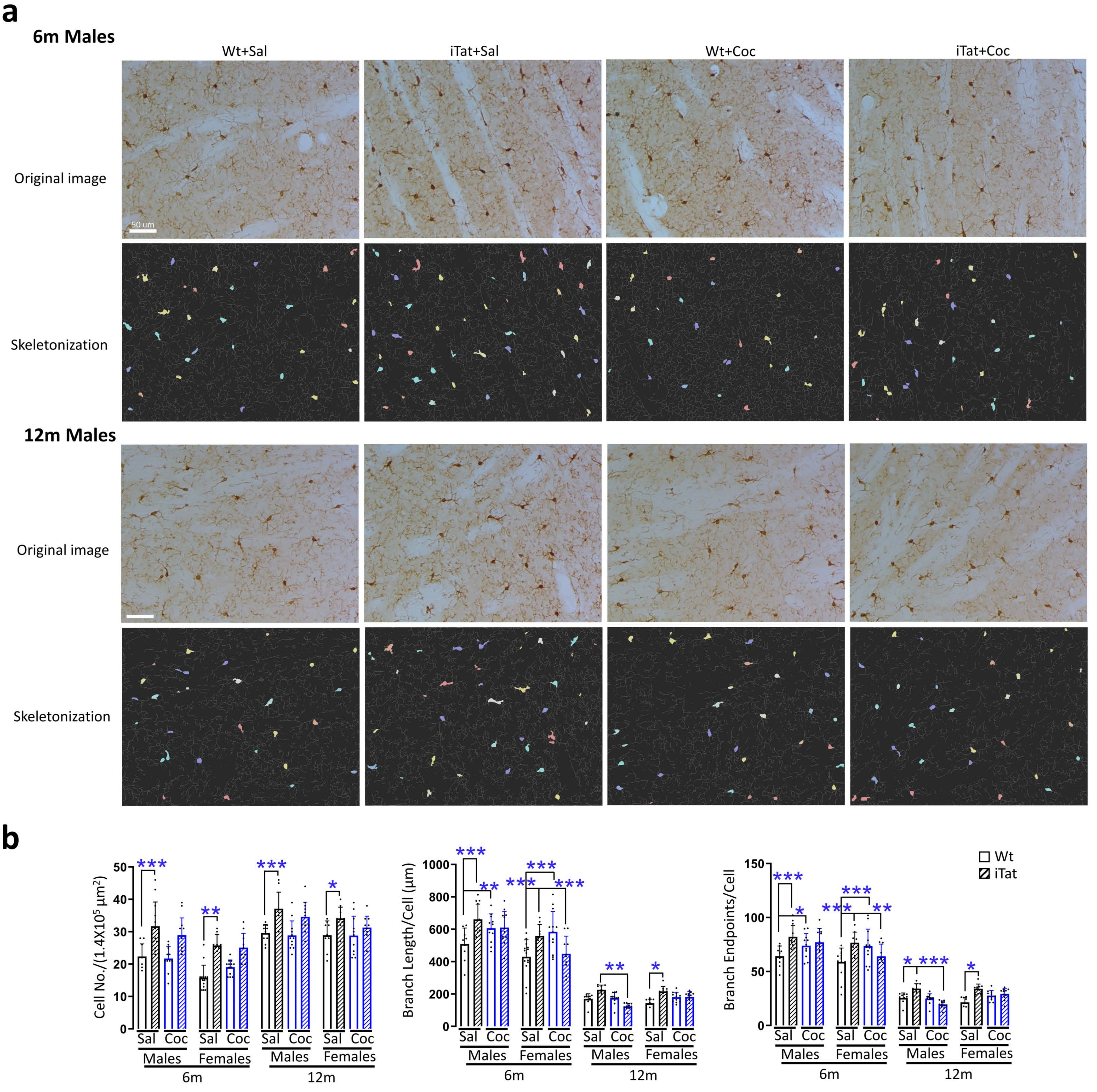

a
6m Males
Wt+Sal
iTat+Sal
Wt+Coc
iTat+Coc
Original image
50 um
Skeletonization
12m Males
Original image
Skeletonization
b
Cell No./(1.4X10$^5$ μm$^2$)
Branch Length/Cell (μm)
Branch Endpoints/Cell
Wt
iTat
Sal
Coc
Males
Females
6m
12m

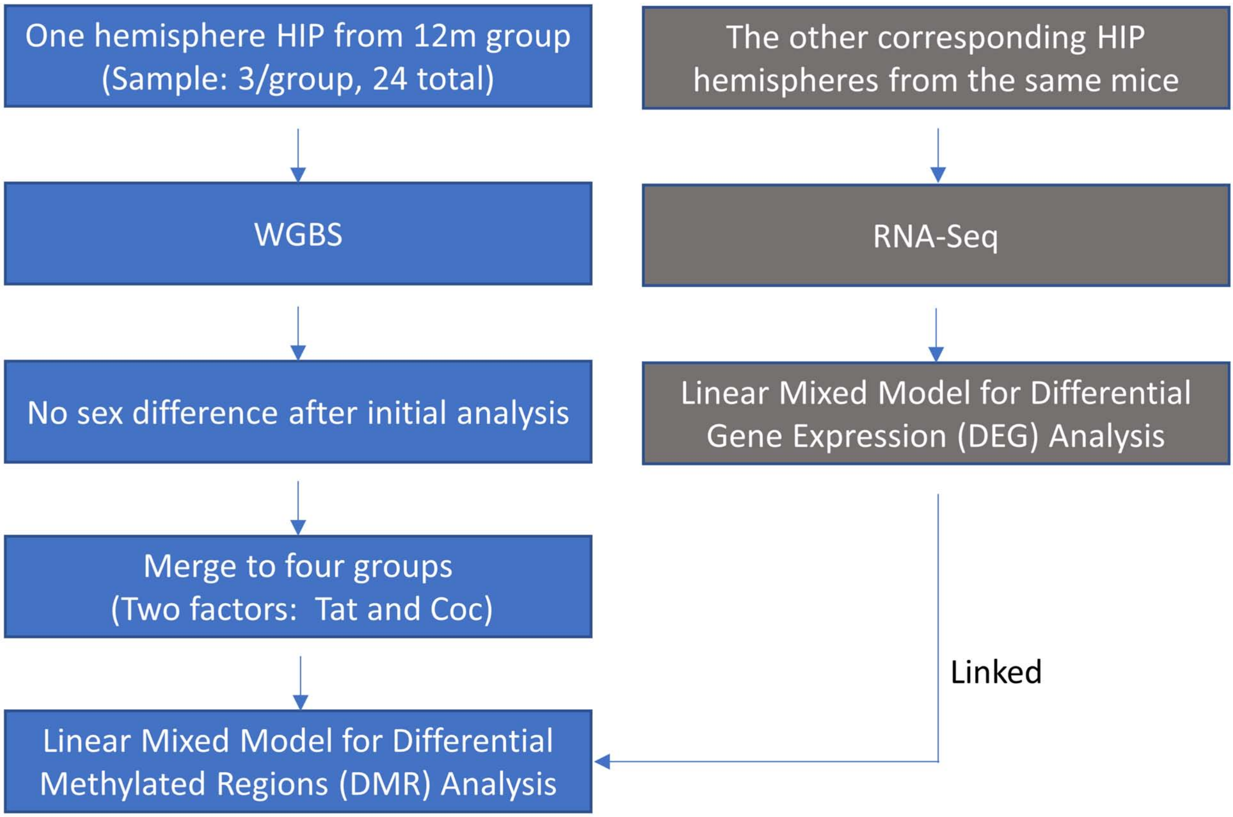
One hemisphere HIP from 12m group (Sample: 3/group, 24 total)
WGBS
No sex difference after initial analysis
Merge to four groups (Two factors: Tat and Coc)
Linear Mixed Model for Differential Methylated Regions (DMR) Analysis
The other corresponding HIP hemispheres from the same mice
RNA-Seq
Linear Mixed Model for Differential Gene Expression (DEG) Analysis
Linked

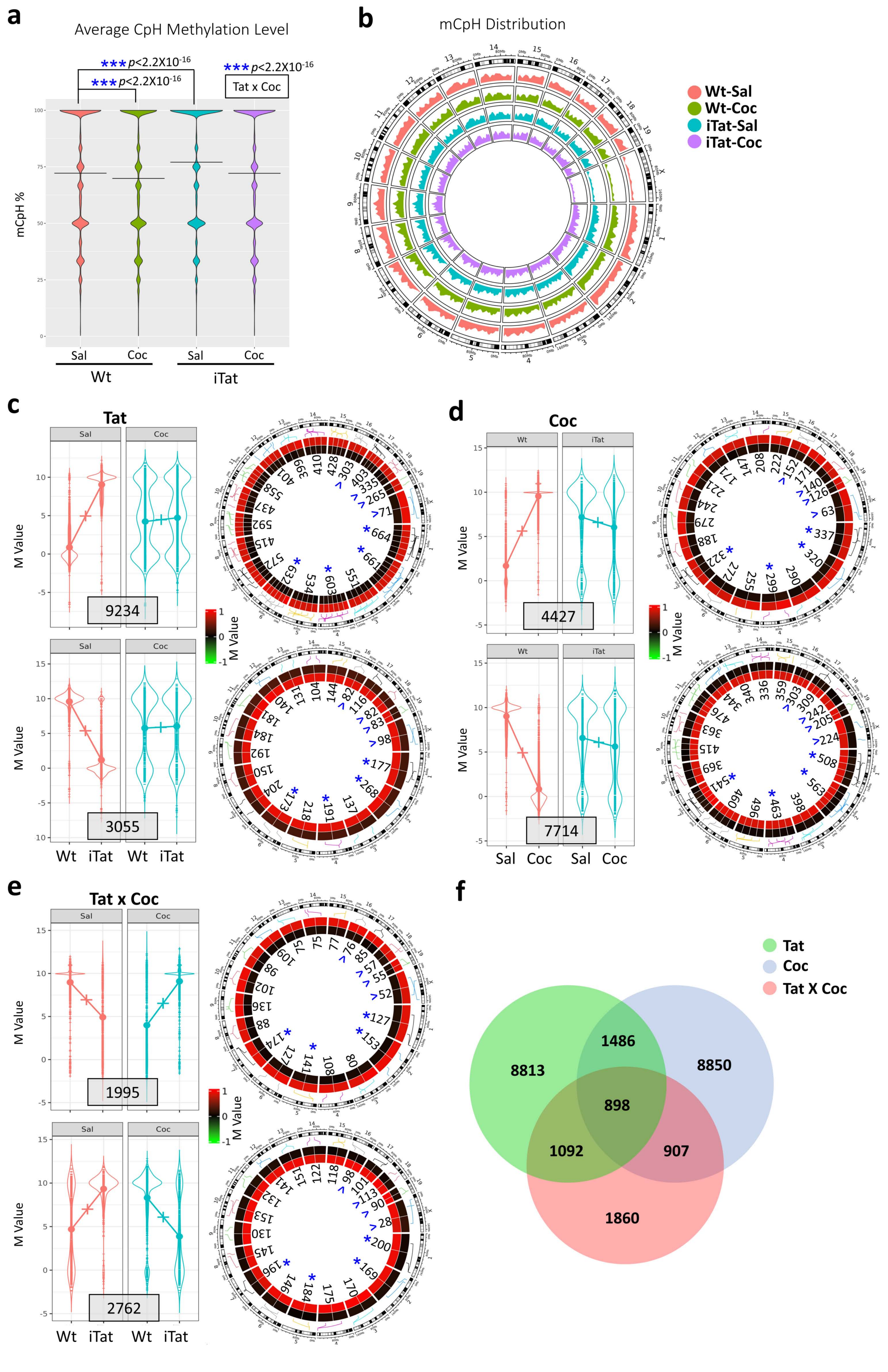

a
Average CpH Methylation Level
*** p<2.2X10^-16
*** p<2.2X10^-16
*** p<2.2X10^-16
Tat x Coc
mCpH %
Sal Coc Sal Coc
Wt iTat
b
mCpH Distribution
Wt-Sal
Wt-Coc
iTat-Sal
iTat-Coc
c
Tat
9234
3055
M Value
Wt iTat Wt iTat
d
Coc
4427
7714
Sal Coc Sal Coc
e
Tat x Coc
1995
2762
f
Tat
Coc
Tat X Coc
8813
1486
8850
898
1092
907
1860

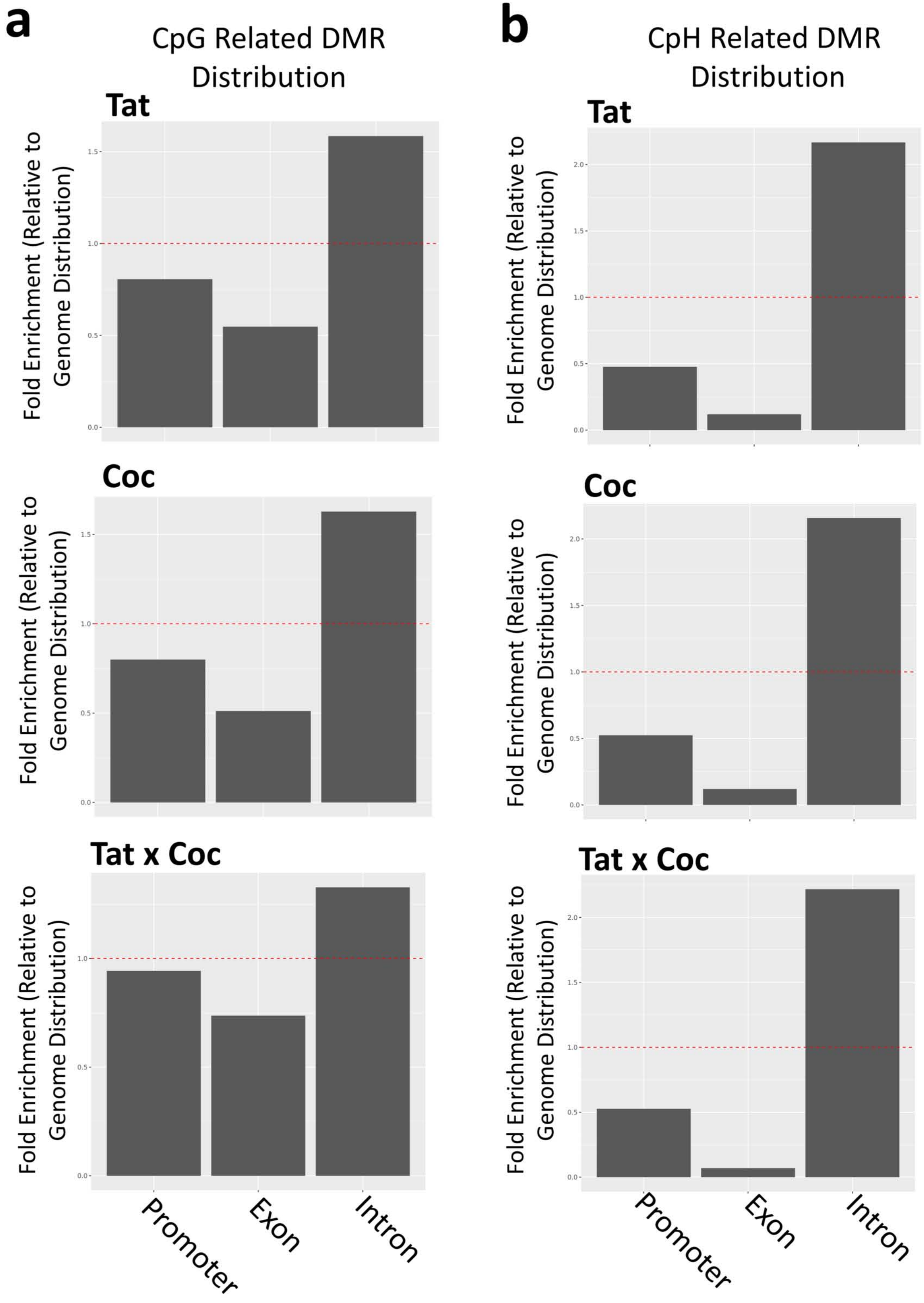
a
CpG Related DMR Distribution
b
CpH Related DMR Distribution
Tat
Coc
Tat x Coc
Fold Enrichment (Relative to Genome Distribution)
Promoter
Exon
Intron

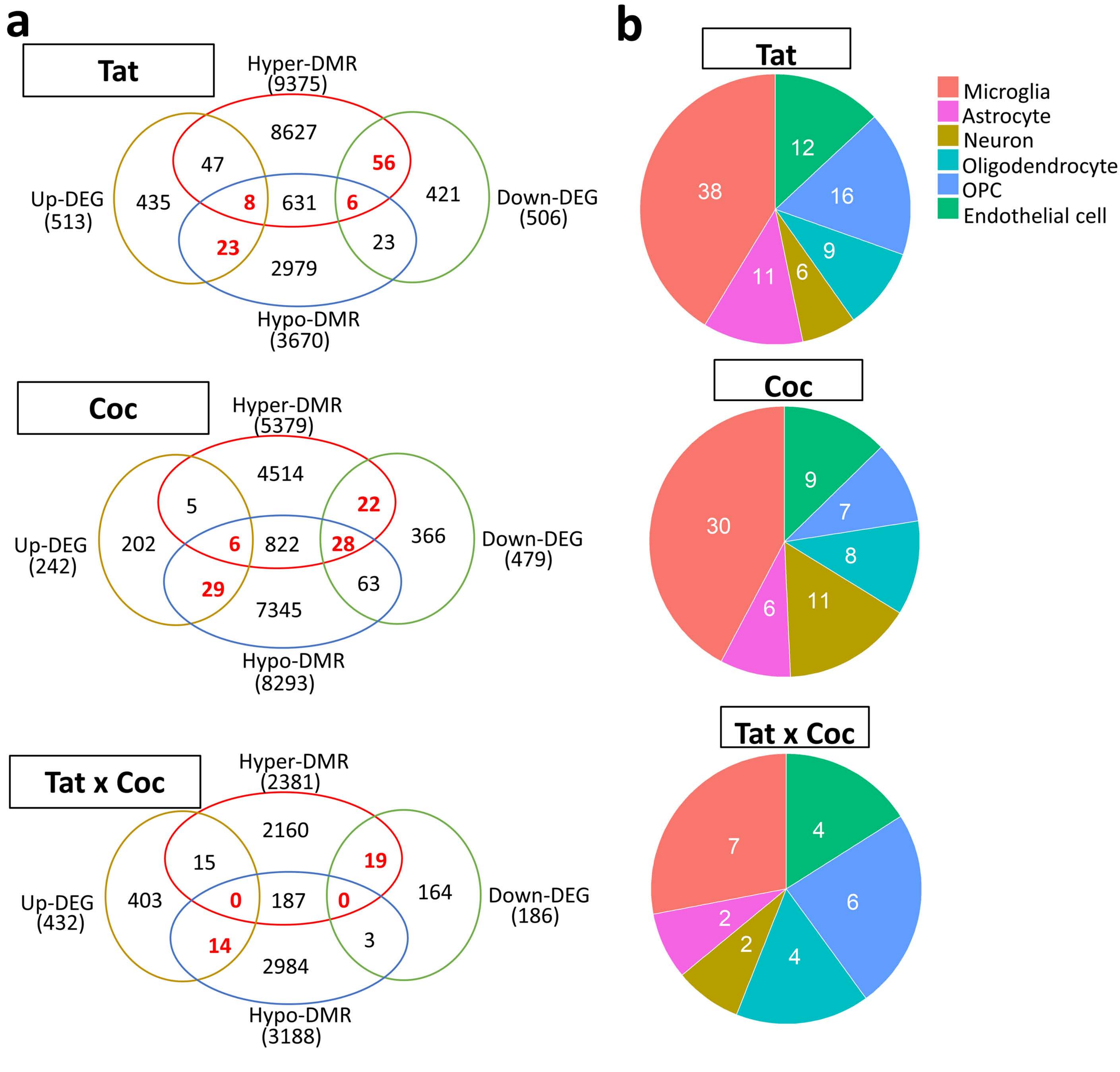
a
Tat
Hyper-DMR
(9375)
8627
47
56
Up-DEG
(513)
435
8
631
6
421
Down-DEG
(506)
23
23
2979
Hypo-DMR
(3670)
Coc
Hyper-DMR
(5379)
4514
5
22
Up-DEG
(242)
202
6
822
28
366
Down-DEG
(479)
29
63
7345
Hypo-DMR
(8293)
Tat x Coc
Hyper-DMR
(2381)
2160
15
19
Up-DEG
(432)
403
0
187
0
164
Down-DEG
(186)
14
3
2984
Hypo-DMR
(3188)
b
Tat
Microglia
Astrocyte
Neuron
Oligodendrocyte
OPC
Endothelial cell
38
12
16
9
6
11
Coc
30
9
7
8
11
6
Tat x Coc
7
4
6
4
2
2

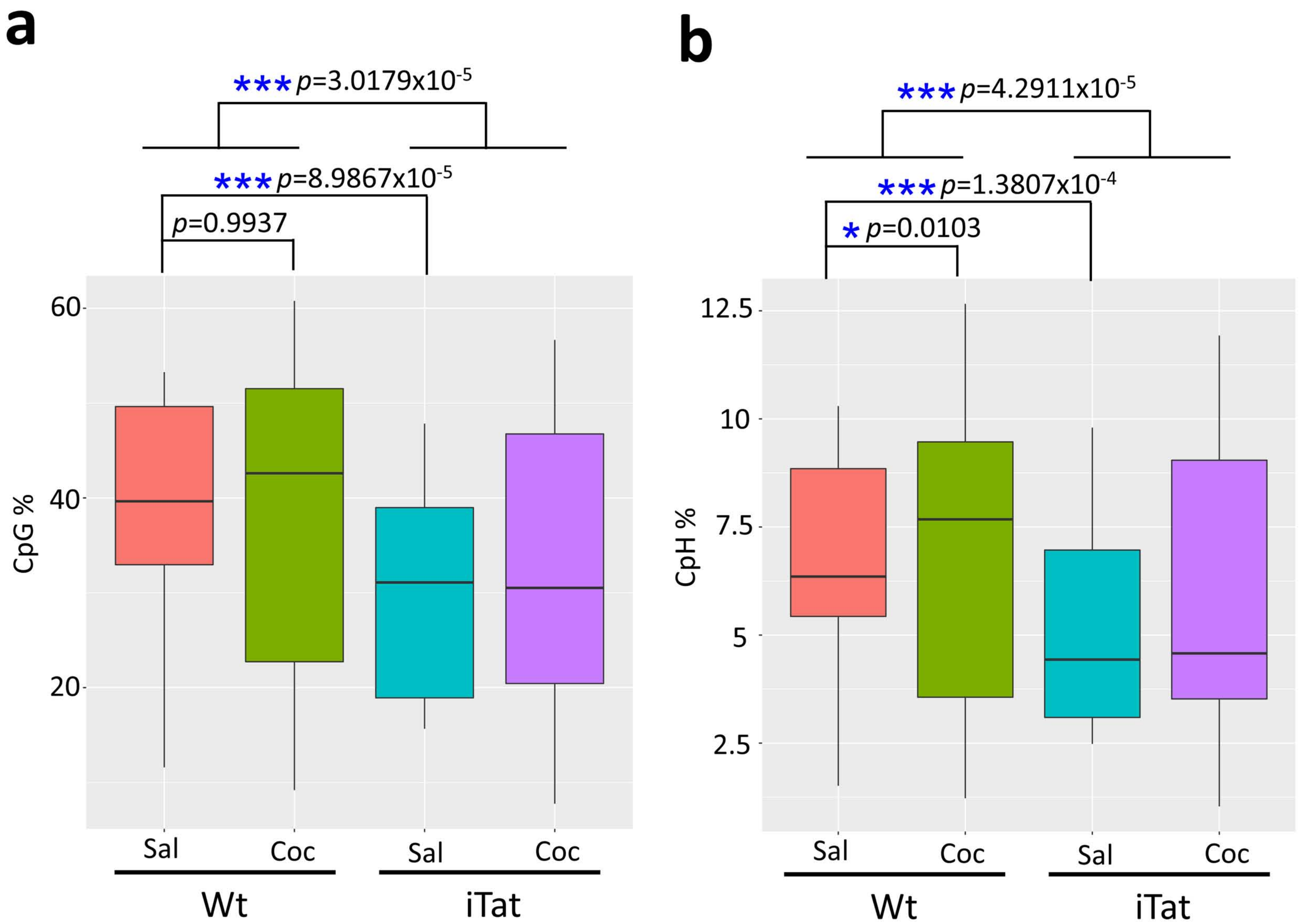
a
***p=3.0179x10-5
***p=8.9867x10-5
p=0.9937
CpG %
60
40
20
Sal
Coc
Sal
Coc
Wt
iTat
b
***p=4.2911x10-5
***p=1.3807x10-4
*p=0.0103
CpH %
12.5
10
7.5
5
2.5
Sal
Coc
Sal
Coc
Wt
iTat

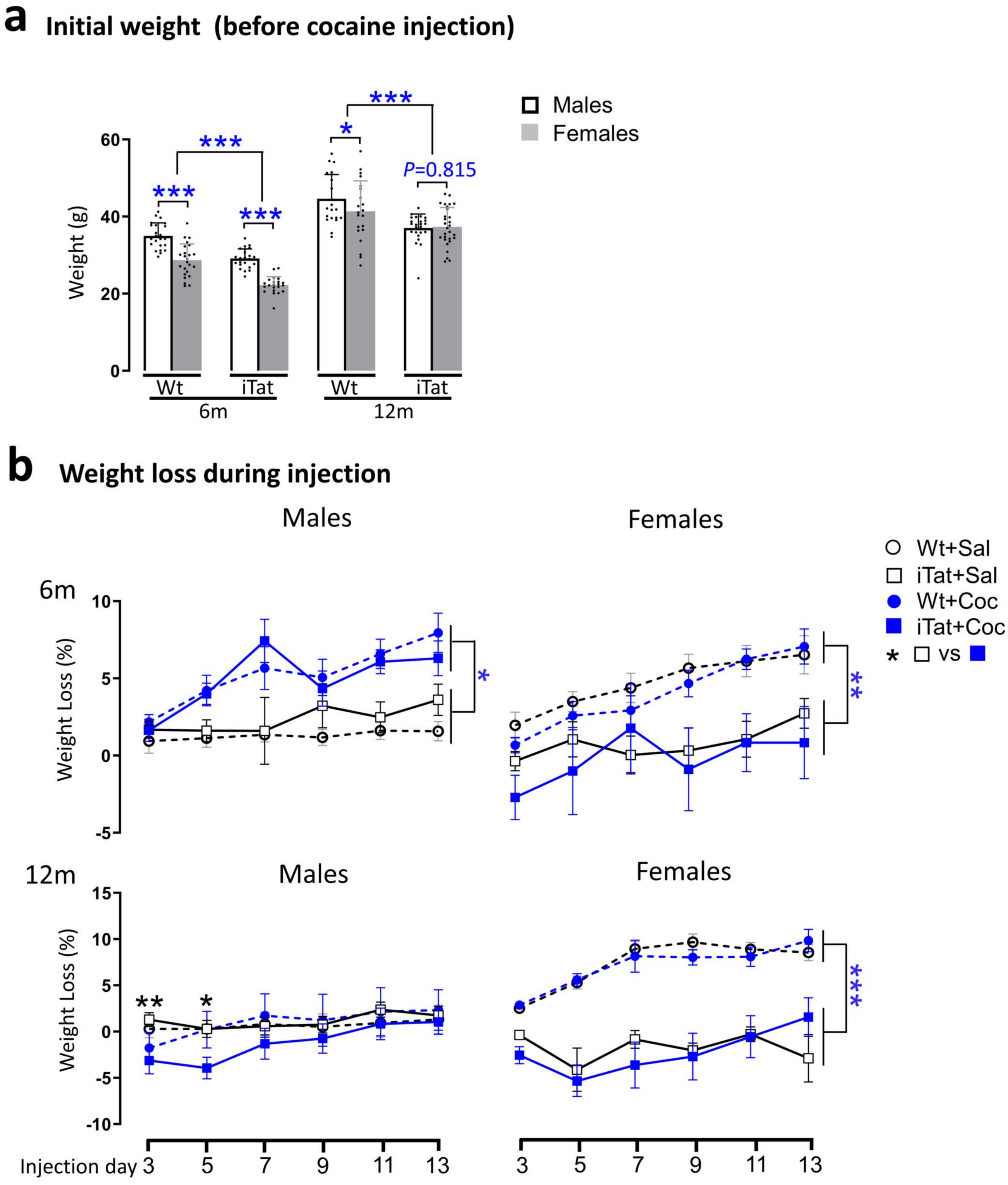

a
Initial weight (before cocaine injection)
Males
Females
Weight (g)
60
40
20
0
***
***
***
*
***
P=0.815
Wt
iTat
Wt
iTat
6m
12m
b
Weight loss during injection
Males
Females
Wt+Sal
iTat+Sal
Wt+Coc
iTat+Coc
* □ vs ■
6m
Weight Loss (%)
10
5
0
-5
*
**
12m
Males
Females
15
10
5
0
-5
-10
**
*
***
Injection day 3
5
7
9
11
13
3
5
7
9
11
13

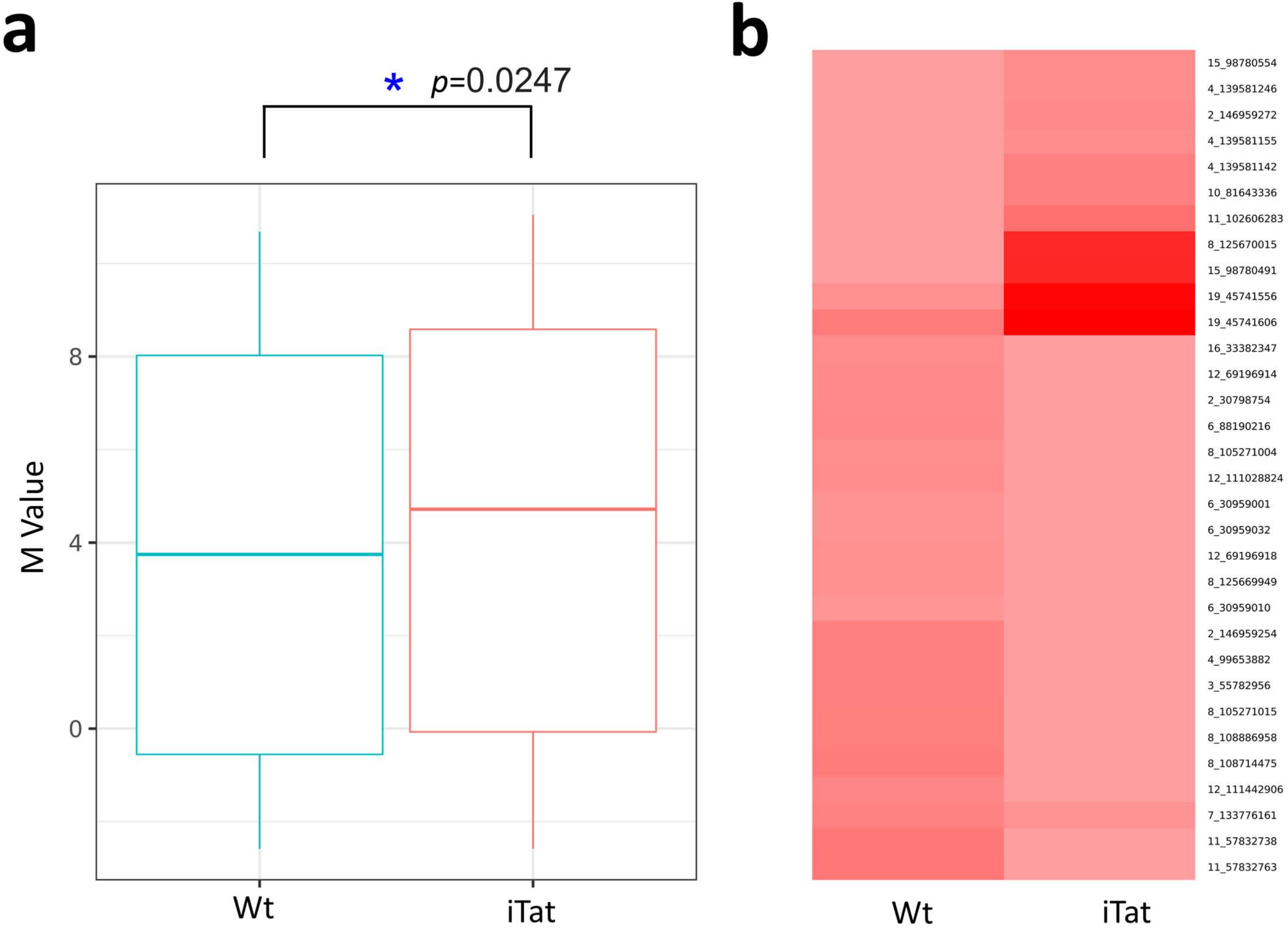

a
b
* p=0.0247
M Value
8
4
0
Wt
iTat
15_98780554
4_139581246
2_146959272
4_139581155
4_139581142
10_81643336
11_102606283
8_125670015
15_98780491
19_45741556
19_45741606
16_33382347
12_69196914
2_30798754
6_88190216
8_105271004
12_111028824
6_30959001
6_30959032
12_69196918
8_125669949
6_30959010
2_146959254
4_99653882
3_55782956
8_105271015
8_108886958
8_108714475
12_111442906
7_133776161
11_57832738
11_57832763
Wt
iTat
